# DDR1-INDUCED NEUTROPHIL EXTRACELLULAR TRAPS DRIVE PANCREATIC CANCER METASTASIS

**DOI:** 10.1101/2020.08.05.238097

**Authors:** Jenying Deng, Yaan Kang, Chien-Chia Cheng, Xinqun Li, Bingbing Dai, Matthew H. Katz, Taoyan Men, Michael P. Kim, Eugene A. Koay, Huocong Huang, Rolf A. Brekken, Jason B. Fleming

**Affiliations:** Department of Surgical Oncology, The University of Texas MD Anderson Cancer Center, Houston, Texas; Department of Molecular and Cellular Oncology, The University of Texas MD Anderson Cancer Center, Houston, Texas; Functional Genomics Core, The University of Texas MD Anderson Cancer Center, Houston, Texas; Genetics, Division of Basic Science Research, The University of Texas MD Anderson Cancer Center, Houston, Texas; Department of Radiation Oncology, The University of Texas MD Anderson Cancer Center, Houston, Texas; Department of Surgery and Hamon Center for Therapeutic Oncology Research, Division of Surgical Oncology, The University of Texas Southwestern Medical Center, Dallas, Texas; Department of Pharmacology, The University of Texas Southwestern Medical Center, Dallas, Texas; Department of Gastrointestinal Oncology, H. Lee Moffitt Cancer Center, Tampa, Florida

**Author notes:** Corresponding Author: Jason B. Fleming, MD.

**Keywords:** DDR1, neutrophil extracellular traps, pancreatic cancer, metastasis, microenvironment

## Abstract

Pancreatic ductal adenocarcinoma (PDAC) tumors are characterized by a desmoplastic reaction and dense collagen that is known to promote cancer progression. A central mediator of pro-tumorigenic collagen signaling is the receptor tyrosine kinase discoid domain receptor 1 (DDR1). DDR1 is a critical driver of a mesenchymal and invasive cancer cell PDAC phenotype. Previous studies have demonstrated that genetic or pharmacologic inhibition of DDR1 prevents PDAC tumorigenesis and metastasis. Here, we investigated whether DDR1 signaling has cancer cell non-autonomous effects that promote PDAC progression and metastasis. We demonstrate that collagen-induced DDR1 activation in cancer cells is a major stimulus for CXCL5 production, resulting in the recruitment of tumor-associated neutrophils (TANs), the formation of neutrophil extracellular traps (NETs) and subsequent cancer cell invasion and metastasis. Moreover, we have identified that collagen-induced CXCL5 production was mediated by a DDR1-PKCθ-SYK-NFκB signaling cascade. Together, these results highlight the critical contribution of collagen I-DDR1 interaction in the formation of an immune microenvironment that promotes PDAC metastasis.

**Summary:** Deng et al find that collagen signaling via DDR1 on human pancreatic cancer cells drives production and release of the cytokine, CXCL5, into systemic circulation. CXCL5 then triggers infiltration of neutrophils into the tumor where they promote cancer cell progression.

## Introduction

Pancreatic ductal adenocarcinoma (PDAC) is now the third-leading cause of cancer death in the United States. The majority of PDAC patients are found to have metastatic disease at diagnosis, and only ∼10% of patients survive over 5 years ^1,2^. One of the major contributors to the dismal prognosis of PDAC is its unique stroma. PDAC is characterized by a desmoplastic reaction accompanying the progression of the disease, resulting in the deposition of a dense extracellular matrix (ECM) ^3^. One of the major ECM components is collagen, which can promote cancer cell survival and facilitate invasion ^4,5^. However, the mechanisms through which collagen functions to promote PDAC progression are not clear. The discoid domain receptor (DDR) family, which includes DDR1 and DDR2, is the only receptor tyrosine kinase family that is specifically activated by fibrillar collagens ^5,6^. Upon ligation with collagen, DDRs undergo auto-phosphorylation and propagate downstream signaling. DDRs have been shown to regulate cancer cell survival, adhesion, proliferation, motility, and invasion in different cancer settings ^7^. In PDAC, elevated expression of DDR1 is negatively associated with clinical outcomes ^8^. In addition, collagen-DDR1 signaling can induce an invasive phenotype in pancreatic cancer cells through an epithelial-mesenchymal transition (EMT) ^9,10^. Therefore, DDR1 might be a critical mediator of collagen-driven tumorigenesis in PDAC. This is supported by a recent study involving genetically ablated DDR1 in a genetically engineered mouse model (GEMM) of PDAC ^11^. As a result, the progression of tumors was significantly delayed and the tumors failed to progress into an undifferentiated phenotype. Moreover, the metastases of the DDR1-deficient tumors were also significantly reduced. We reported in an earlier study that the pharmacological inhibition of DDR1 activation by a novel small-molecule inhibitor, 7rh benzamide, inhibited tumorigenesis and enhanced chemosensitivity in orthotopic xenografts and autochthonous pancreatic tumors ^12^.

Although DDR1 induces an invasive cancer cell phenotype that contributes to invasion, metastasis, and therapy resistance, whether DDR1 can mediate a communication between cancer cells and stromal cells and alter the tumor microenvironment (TME) is poorly understood. This is particularly important because metastasis is a complex, multistep process requiring cancer cell migration and survival in a distant organ ^13^. Therefore, a TME that facilitates the distant travel and seeding of cancer cells is essential. The recruitment of immunosuppressive stromal cells, including tumor-associated macrophages (TAMs) and tumor-associated neutrophils (TANs), by cancer cells is a major contributor to a metastasis-permissive TME ^14^. As a critical component of the innate immune system, neutrophils are first responders to infection and injury ^15^. Through the generation of reactive oxidants, and the activation of granular constituents and neutrophil extracellular traps (NETs), neutrophils target microbes and prevent their dissemination ^16^. In cancer, there is growing evidence that TANs can enhance tumor progression through NETs ^17^. NETs are long, thin-stranded, web-like extracellular fibers formed by neutrophils, consisting of chromatin DNA filaments and specific proteins, such as lactoferrin, myeloperoxidase (MPO), histones, and neutrophil elastase. Recent studies have shown that NETs are highly associated with metastasis in different cancer types ^18-22^. NETs can remodel the stroma to help tumor cells invade and induce thrombosis formation to help tumor cell clusters metastasize ^21,23^. NETs also capture and induce apoptosis in cytotoxic T cells ^24^. Moreover, a recent study has shown that NETs can activate CCDC25 on cancer cells and enhance cell motility ^22^. In spite of the important functions of TANs and NETs during cancer metastasis, the pathways that cancer cells use to induce NET formation remain unclear.

In this study, we investigated whether cancer cells exploit collagen-DDR1 signaling to communicate with other stromal cells and modulate the TME to promote PDAC progression and metastasis. The results demonstrated that the activation of cancer cell DDR1 by collagen is an essential step for TAN infiltration and NET formation. Specifically, we identified CXCL5 as a key chemokine in collagen I-induced NET formation and demonstrated that the pharmacologic blockade of DDR1 effectively prevents collagen induction of CXCL5, subsequent NET formation, and cancer cell invasion. Moreover, we also identified that the collagen-induced CXCL5 production was mediated by a PKCθ-SYK-NFκB signaling cascade downstream of activated DDR1. Taken together, we conclude that collagen stimulates pancreatic cancer cells to produce CXCL5 through a DDR1-PKCθ-SYK-NFκB pathway, and as a result, CXCL5 induces TANs to form NETs and promote cancer cell invasion and metastasis.

## Results

### DDR1 drives metastasis in PDAC

Recent evidence demonstrates that the genetic ablation of DDR1 in a GEMM of PDAC resulted in a significant reduction of metastasis ^11^. To further validate this result, we exploited widely used human PDAC cell lines, human PDAC cell lines generated in our lab (MDA-PATC), and patient-derived xenograft (PDX) tumors. We first examined the expression of DDRs in PDAC cell lines and confirmed that 14 of the cell lines expressed high levels of DDR1 while only several expressed DDR2 (Figure 1A). Then we orthotopically implanted two cell lines with robust DDR1 expression, PATC 148 and PATC 153, into nude mice and harvested liver metastases which were used to generate matched metastatic cell lines (PATC 148LM and 153LM). We compared MDA-PATC 148LM and 153LM cells with the matched parental lines and found that the metastatic clones expressed higher levels of DDR1 (Figure 1A). Comparison of PDX tumors derived from metastatic or primary human PDAC tumors revealed that PDX derived from metastatic tumors expressed higher levels of DDR1 than PDX derived from primary tumors (Figure 1B).

**Figure 1.**
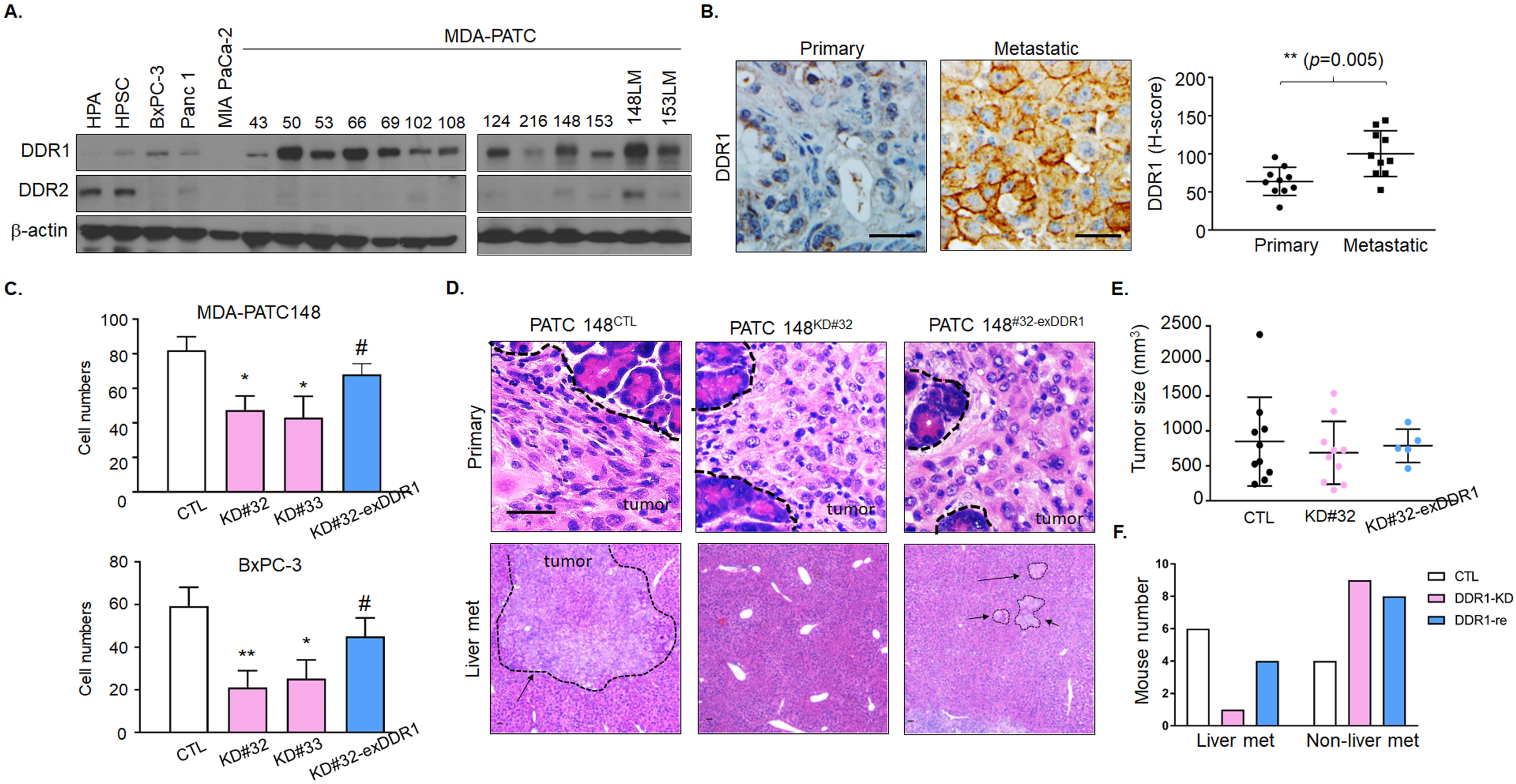
DDR1 Induces liver metastasis in pancreatic cancer. **A** DDR1 and DDR2 expression were analyzed by western blotting in 2 fibroblasts, 14 primary PDAC cell lines and 2 of metastatic PDAC cell lines. **B** DDR1 were observed at PDX tumors derived from metastatic or primary human PDAC tumors by immunohistochemical staining using anti- human DDR1 antibody and identified using PE Vectra3. Scale bar, 50 μm. The H-score of DDR1 quantification was displayed as DBA signals by inform software. **C** Cell invasion assay in MDA-PATC 148 cells with knockdown or re-expression DDR1 were used by matrigel transwell chamber. The invading cells in each chamber were counted under a fluorescence microscope after culture 18 hours, and the average number of cells was calculated based on the number of cells found in six fields per chamber. **D**, **E**, **F** Mice were orthotopically injected with MDA-PATC 148 (control, DDR1-deficient or DDR1-reexpression clones) cells for 9 weeks. **D** H&E staining of pancreas and liver section. *Arrow*: region of tumor (10 mice per group), Scale bar, 50 μm. **E** Tumors size measurement in pancreas. **F** The numbers of liver-met calculation (*n* = 10 for each group).

To confirm DDR1 induced cancer cell invasion and metastasis, we generated stable DDR1-deficient clones using different shRNAs against DDR1 in MDA-PATC 148 (MDA-PATC 148^KD#32^ and MDA-PATC 148^KD#33^) and BxPC-3 (BxPC-3^KD#32^ and BxPC-3^KD#33^) cells (Supplementary Figure 1A). In addition, we rescued DDR1 expression by introducing an shRNA-resistant DDR1 construct in MDA-PATC 148^KD#32^ (MDA-PATC 148^KD#32-exDDR1^) and BxPC-3^KD#32^ (BxPC-3^#32-exDDR1^) cells (Supplementary Figure 1B). The loss of DDR1 resulted in a reduction of invading cells in each cell line and this effect was rescued by DDR1 re-expression (Figure 1C). Upon orthotopic implantation of MDA-PATC 148^CTL^, MDA-PATC 148^KD#32^, and MDA-PATC 148^KD#32-exDDR1^ cells, we found that DDR1 knockdown in the cancer cells had no effect on primary tumor growth, but resulted in a significant reduction of liver metastasis (WT, 80%; KD^#32^, 20%; *p* value = 0.036 in Fisher’s exact test) (Figure 1D, 1E and 1F).

### DDR1 induces CXCL5 production and Ly6G^+^ neutrophil infiltration

To investigate the influence of cancer cell DDR1 signaling on the TME, we screened for cancer cell DDR1-induced cytokine production using a human chemokine antibody array. We found that the activation of DDR1 by collagen induced the production of four candidate factors, CD130, CXCL8, CXCL5, and MCP-1, which were reduced by knocking down DDR1 (Figure 2A). We then validated that CXCL5 was a DDR1-induced factor by real-time qPCR assays. (Supplementary Figure 2). The exposure of parental cancer cells to collagen I for 3 h increased mRNA and protein levels of CXCL5 in MDA-PATC 148 and BxPC-3 cells, which was diminished in knockdown DDR1 cells (Figure 2B and 2C). In addition, the mRNA and protein expression of CXCL5 was rescued after the re-expression of DDR1 in MDA-PATC 148^KD#32^ and BxPC-3^KD#32^ cells (Figure 2B and 2C). To further confirm this was a DDR1-mediated effect, we also overexpressed DDR1 in an additional five human pancreatic cancer cell lines, which resulted in a significant increase of CXCL5 expression upon collagen activation (Figure 2D).

**Figure 2.**
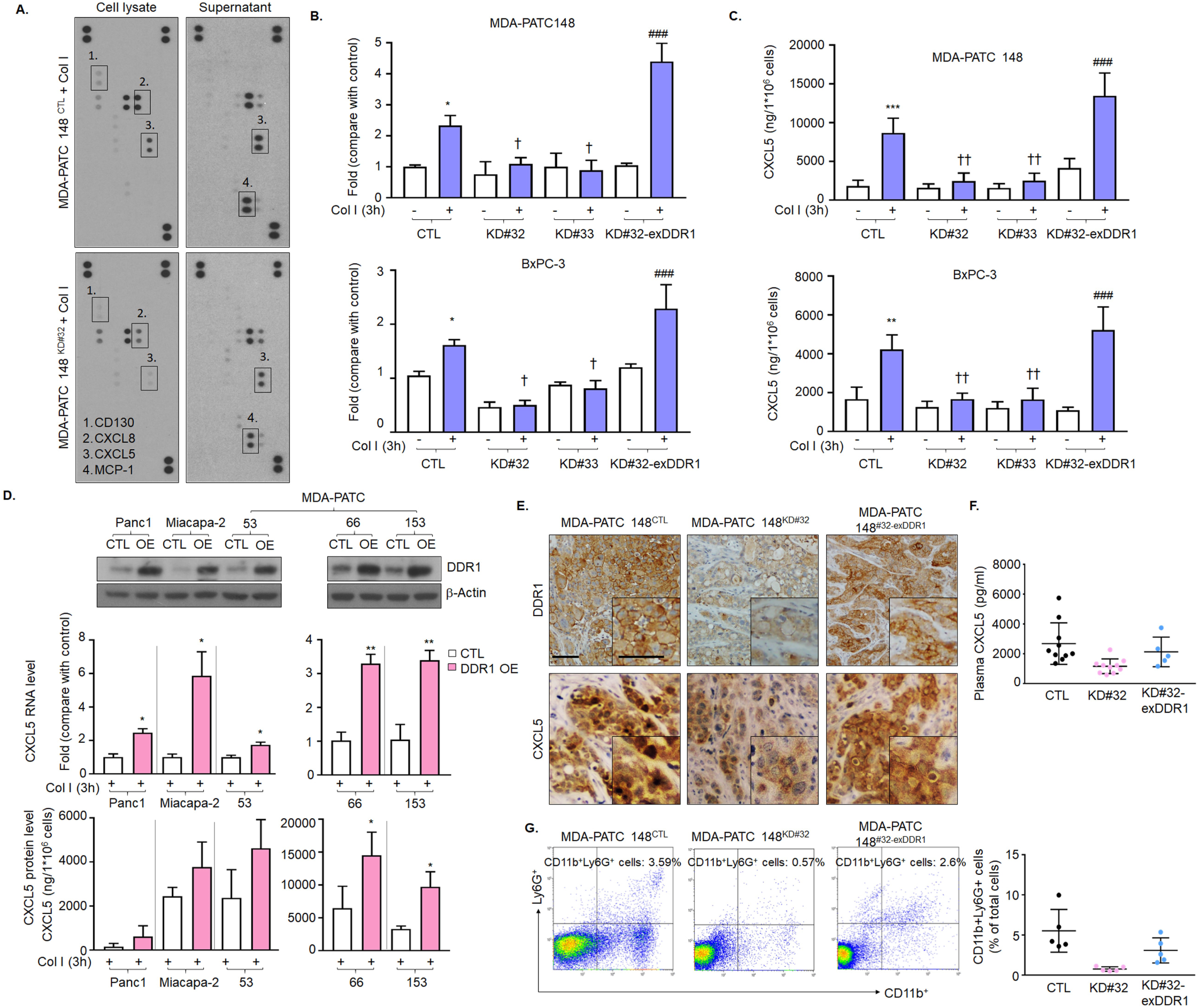
DDR1 Induces CXCL5 production in pancreatic cancer cells. **A** chemokine array analysis in cell lysate and supernatant of MDA-PATC 148 cells with knockdown DDR1. **B and C** MDA-PATC 148 cells with knockdown or re-express DDR1 were treated with collagen I for 3 hours. **B** CXCL5 mRNA level by using real-time PCR. **C** Protein level by using ELISA. **D** overexpressed DDR1 in 5 pancreatic cancer cell lines, *upper:* DDR1 level were checked by western; *middle:* CXCL5 level were detected by real-time PCR; *bottom:*. CXCL5 level were analyzed by ELISA. **E**, **F and G** Mice were orthotopically injected with MDA-PATC 148 (control, DDR1-deficient or DDR1-reexpression clones) cells for 9 weeks. **E** Immunohistochemical staining with anti-DDR1 (upper panel) and anti-CXCL5 (bottom panel) antibodies in pancreas (*n*= 10 for each group). **F** ELISA showed CXCL5 level in plasma harvest from mice **(*n* =** 10 for each group). **G** *Left:* FACS by using anti-CD11b and anti-Ly6G antibodies to determine the presence of CD11b^+^Ly6G^+^ neutrophils infiltration in pancreas; *right*. The data were collected from three independent experiments **(*n*** = 5 for each group).

CXCL5 is also known as epithelial-derived neutrophil-activating peptide 78 (ENA-78) and has been previously shown to induce TAN infiltration and increase the metastatic risk in hepatocellular carcinoma^25^. We sought to investigate whether DDR1-derived CXCL5 is associated with TAN infiltration in PDAC. We first measured the level of CXCL5 and Ly6G^+^ neutrophil infiltration in tumors derived from MDA-PATC 148 variants (MDA-PATC 148^CTL^, MDA-PATC 148^KD#32^, and MDA-PATC 148^KD#32-exDDR1^). Knocking down DDR1 reduced the level of CXCL5 in the primary tumor and plasma (Figure 2E and 2F). Importantly, we also observed a reduction of CD11b^+^Ly6G^+^ neutrophil infiltration in DDR1-knockdown tumors (Figure 2G). Re-expression of DDR1 (MDA-PATC 148^KD#32^-ex-DDR1) rescued CXCL5 levels in the tumor and plasma and CD11b^+^Ly6G^+^ TAN infiltration (Figure 2E, 2F and 2G). To verify this association in a more clinically relevant setting, we performed immunohistochemistry on a tissue microarray of pancreatic cancer PDXs. Scoring from 82 tumor samples identified a positive correlation between DDR1 expression and CXCL5 production (Pearson *r* = 0.4460; 95% CI, 0.2535-0.6045) and Ly6G^+^ TAN infiltration (*r* = 0.2840; 95% CI, 0.07144-0.4720) (Figure 3A, 3B and 3C). As expected, a positive correlation was also observed between CXCL5 and the infiltration of Ly6G^+^ TANs (*r* = 0.6403; 95% CI, 0.4916-0.7527) (Figure 3A and 3D). To confirm the results were not an artifact of immunodeficient mice necessary for PDX generation, we repeated the experiments in a syngeneic model. KP^wm^C cells (congenic mouse PDAC cells derived from *Kras*^*LSL*.*G12D/+*^; *p53*^*R172H/+*^; *Pdx1*^*CreTg/+*^ [*KP*^*wm*^*C*] mice) with stable DDR1 knockdown were generated (KP^wm^C^KD#588^ and KP^wm^C ^KD#809^) (Supplementary Figure 3A) and orthotopically implanted into immunocompetent C57BL/6J mice. Similar to the xenograft models, DDR1-deficient KP^wm^C orthotopic tumors consistently demonstrated decreased CD45^+^CD11b^+^Ly6G^+^ neutrophil infiltration and fewer liver metastases (WT, 80%; KD^#588^, 0%; KD^#809^, 20%) (Supplementary Figure 3B and C). Together, these results suggest that DDR1 on cancer cells drives CXCL5 production, CD45^+^CD11b^+^Ly6G^+^ TAN infiltration, and liver metastasis.

**Figure 3.**
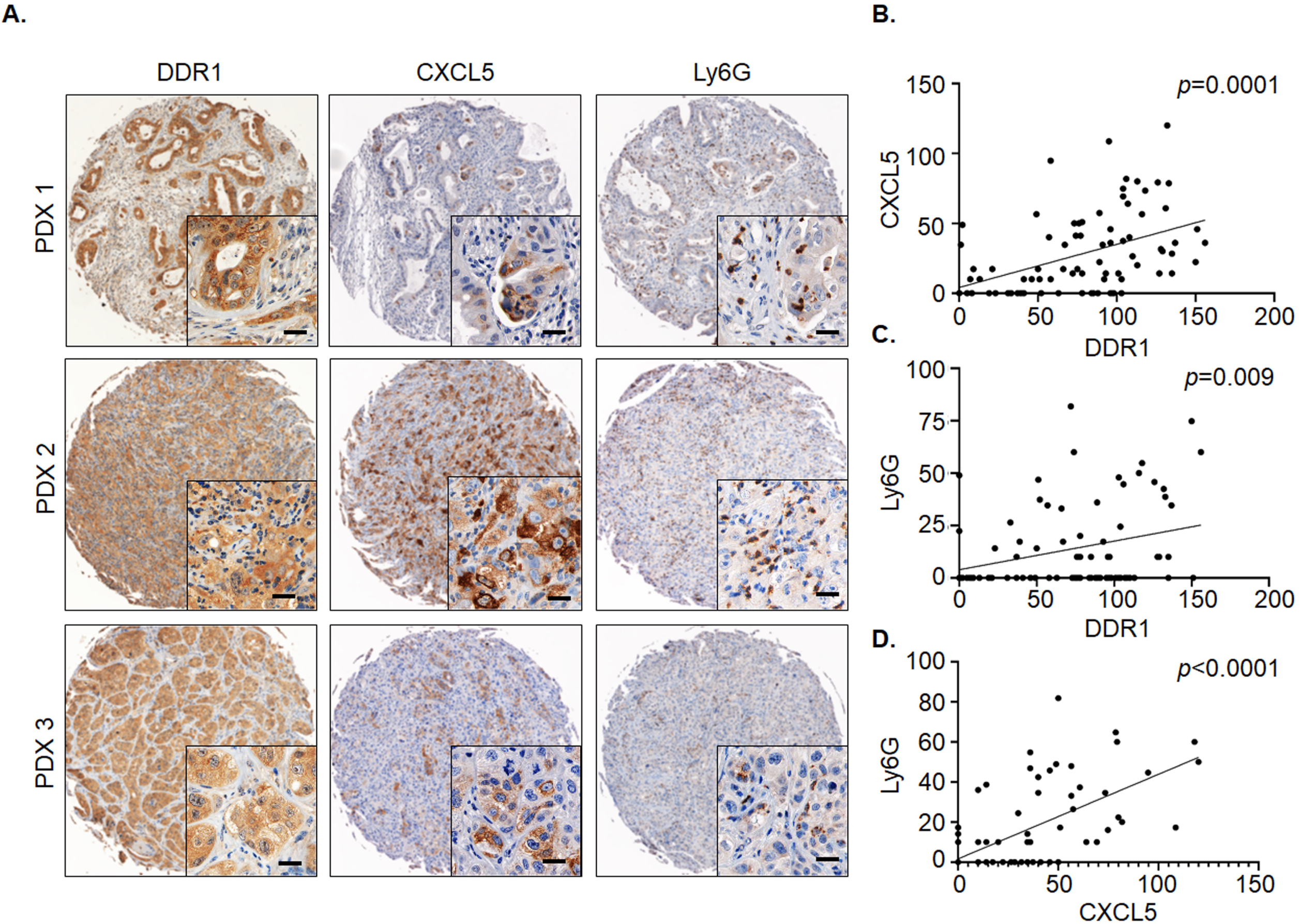
The correlation of DDR1, CXCL5 and neutrophils infiltration at TMA in PDX tumors. **A** immunohistochemical staining showed DDR1, CXCL5 and Ly6G^+^ neutrophils infiltration at PDX tumors and identified using PE Vectra3. Scale bar, 50 μm. **B, C, D** The Pearson correlation showed relationship of DDR1, CXCL5 and Ly6G by using H-score which quantified the DBA signals by inform software.

### DDR1-induced NET formation enhances pancreatic cancer cell invasion

We then asked if the DDR1-induced TAN infiltration directly enhanced the metastatic capability of cancer cells. We first observed a higher level Ly6G^+^ TANs within the TME of tumors derived from cell lines selected for liver metastasis, PATC 148LM and 153LM, when compared with tumors derived from the parental lines (Figure 4A). In addition, PDXs derived from metastatic PDAC tumors were found to have higher numbers of Ly6G^+^ TANs than PDXs derived from primary tumors (Figure 4B).

**Figure 4.**
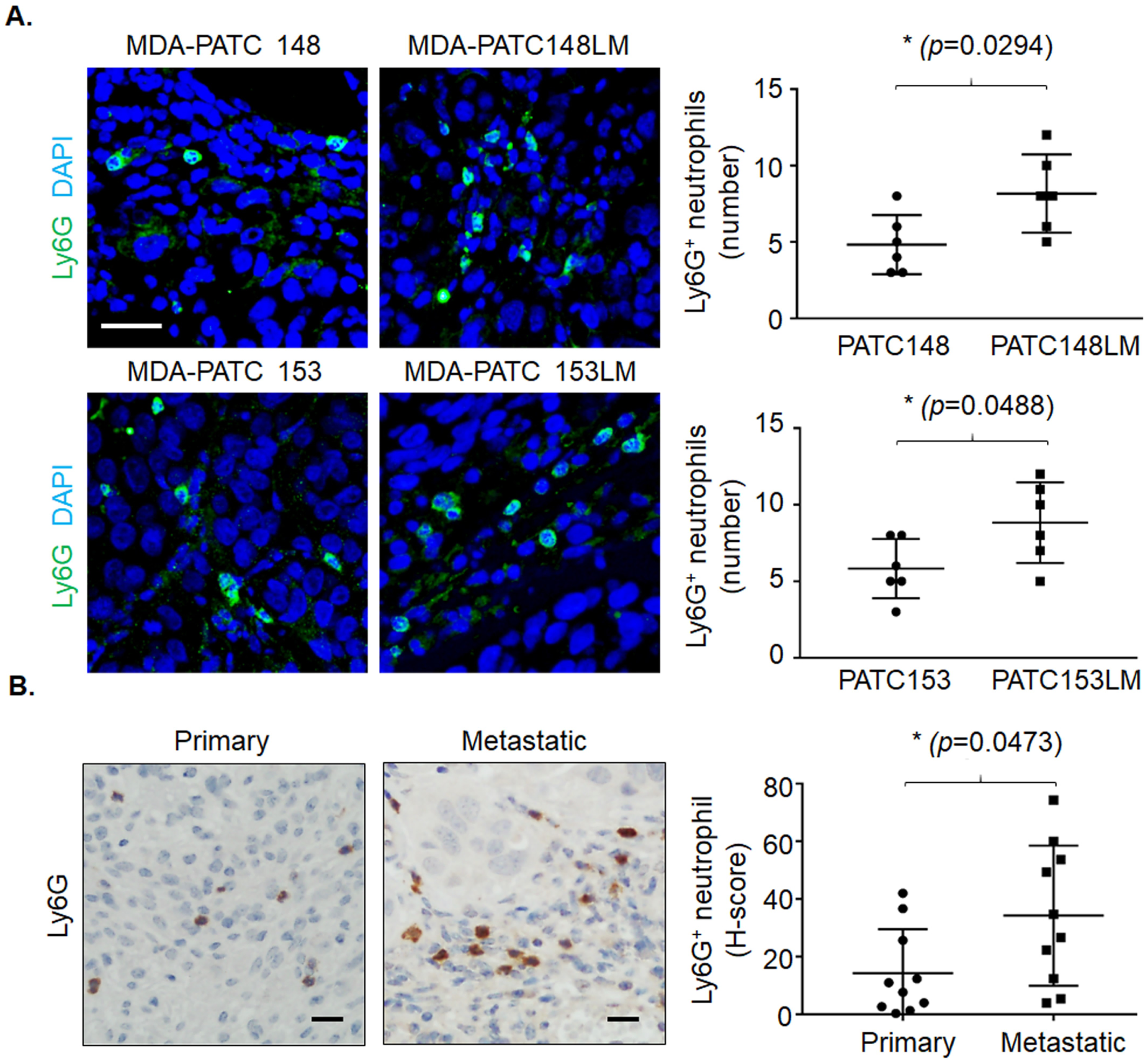
metastatic tumors recruit more Ly6G^+^ neutrophils infiltration than primary tumors. **A** Ly6G^+^ neutrophils were observed at tumors derived from primary and match liver-met cell lines by immunofluorescence staining using anti-Ly6G (green) and DAPI (blue) with a fluorescence microscope. Scale bar, 50 μm. The number of neutrophils were counted in 20X field, 6 fields per slice. **B** Ly6G^+^ neutrophils were observed at PDX tumors derived from metastatic or primary human PDAC tumors by immunohistochemical staining using anti-Ly6G antibody and identified using PE Vectra3. Scale bar, 50 μm. The H-score of Ly6G quantification was displayed as DBA signals by inform software.

Recent studies have demonstrated NET formation is a major driver of metastasis in breast cancer^21^. In PDAC, NETs have also been frequently observed in tumor tissues ^18^. Thus, we investigated whether NETs contributed to DDR1-induced cancer cell invasion. NET formation requires the generation of reactive oxygen species and nicotinamide adenine dinucleotide phosphate (NADPH) oxidase activity as well as the activation of peptidylarginine 4 (PAD4) that promotes the decondensation of nuclear DNA by histone citrullination and the release of MPO/neutrophil elastase (NE) from azurphilic granules ^26,27^. These events can be triggered *in vitro* by exposing neutrophils to phorbol myristate acetate (PMA) ^28^. We co-cultured neutrophils harvested from the blood of patients with untreated PDAC, with MDA-PATC 148 or BxPC-3 cells for 18 h using a Matrigel-coated transwell chamber and found increased NET formation and citrullinated histone H3 expression (Figure 5A, 5B, and 5C). Histone H3 citrullination and NET formation was significantly reduced when the experiments were performed using PDAC cells lacking DDR1 (Figure 5A, 5B, and 5C) and was rescued when DDR1 expression was restored (Figure 5A and 5B). When compared with cancer cells alone, the addition of neutrophils resulted in an almost 3-fold increase of invasion in MDA-PATC 148 and BxPC-3 cells. Knocking down DDR1 in the cancer cells significantly reduced neutrophil-mediated cancer cell invasion (Figure 5D). In contrast, the re-expression of DDR1 recovered the effect of neutrophil-mediated cancer cell invasion in MDA-PATC 148^KD#32-exDDR1^ and BxPC-3^KD#32-exDDR1^ cells (Figure 5D). Interestingly, the inhibition of PDA4 and NE inhibited NET formation and cancer cell invasion in co-culture neutrophils with MDA-PATC 148 and BxPC-3 for 18 h (Figure 5E, 5F, and 5G). However, NADPH oxidase inhibition had no effect on NETs and cancer cell invasion, and DNase I treatment showed only a partial effect compared with control (Figure 5E, 5F, and 5G). Taken together, these data suggest that DDR1 on pancreatic cancer cells mediates the formation of NETs which promotes cancer cell invasion through an NAPDH oxidase-independent pathway.

**Figure 5.**
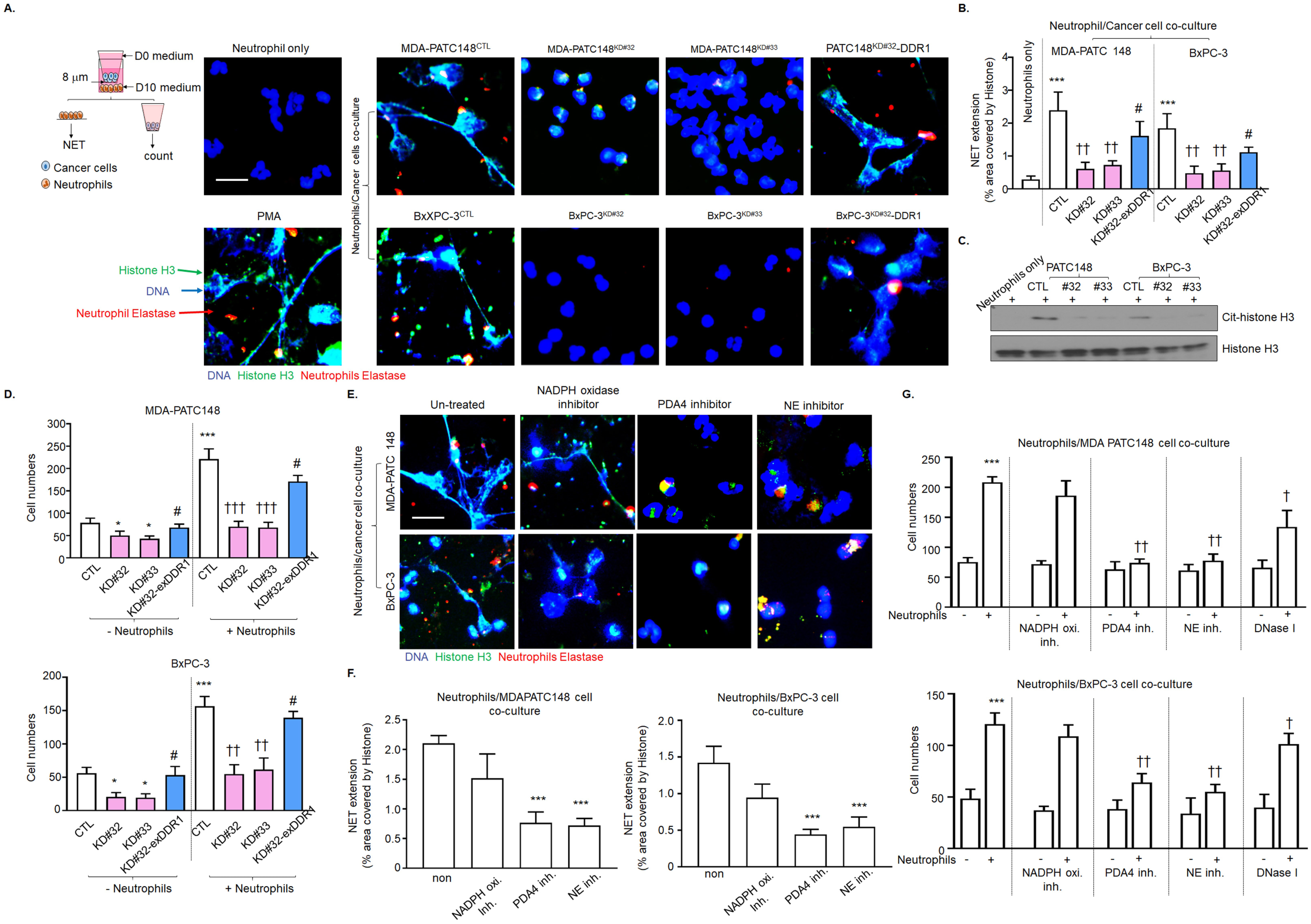
DDR1-positive pancreatic cancer cells mediated NET formation from neutrophils and enhanced cancer cell invasion. **A-G** Human neutrophils were co-culture with DDR1-knockdown or re-expression of MDA-PATC 148 or BxPC-3 by matrigel transwell chamber, with or without NADPH oxidase inhibitor, PDA4 inhibitor, NE inhibitor and Dase I treatment, for 18 hours. **A and E** NET structures were analyzed by immunofluorescence staining using DAPI *(blue)*, anti-NE (*red)* and anti-histone *(green)* mAbs. *Scale bar*, 50 μm. **B and F** The NET quantification is displayed as NET histone area (μm^2^) /per filed. **C** Cit-histone H3 expression were analyzed by western blotting. **D and G** The number of invaded cells analyzed by immunofluorescence staining using DAPI and calculated based on the number of cells found in six fields /per chamber.

### Cancer cell DDR1-induced CXCL5 mediates NET formation and NET-induced cancer cell invasion

To further examine the function of CXCL5 in DDR1-mediated NET formation and cancer cell invasion, the invasion assay was repeated in the presence of a monoclonal CXCL5 neutralizing antibody or recombinant CXCL5. The CXCL5 neutralizing antibody significantly reduced MDA-PATC 148 and BxPC-3 cell-induced histone H3 citrullination, NET formation, and cancer cell invasion (Figure 6A left panel, 6B, 6D and 6F). Conversely, recombinant human CXCL5 induced histone H3 citrullination, NET formation, and cancer cell invasion in MDA-PATC 148^KD #32^ cells (Figure 6A right panel, 6C, 6E, and 6G). The inhibition of PDA4 and NE blocked recombinant CXCL5-mediated, NET-induced MDA-PATC 148^KD#32^ cell invasion, but NADPH inhibition had no effect (Supplementary Figure 4). To evaluate whether the DDR1-induced NET formation and NET-induced cell invasion were mediated by the secretion of soluble factors, we collected conditioned media from three *in vitro* cell culture conditions: cancer cells alone (CCM), neutrophils alone (NCM) or neutrophils exposed to cancer cell conditioned media (NCCM). NET formation and citrullinated histone H3 was induced by CCM from DDR1-expressing cancer cells with collagen I but not from CCM harvested from DDR1-knockdown cancer cells with collagen I stimulation for 3 h (Supplementary Figure 5A, B, and C). We then repeated the cancer cell invasion assay experiment using NCCM in the presence or absence of collagen I. NCCM from MDA-PATC 148^CTL^/collagen I/neutrophil cultures induced cancer cell invasion in a DDR1-independent manner (Supplementary Figure 5D). However, NCCM harvested from MDA-PATC 148^KD#32^/collagen I/neutrophil cultures failed to increase cancer cell invasion again, regardless of the DDR1 status (Supplementary Figure 5E). These carefully designed studies demonstrate that CXCL5 is a soluble factor released into the media from DDR1-positive cancer cells in the presence of collagen; when exposed to CXCL5, neutrophils generate NETs that promote cancer cell invasion. The dependence upon DDR1 is demonstrated by the fact that CCM from DDR1-expressing cancer cell/collagen I cultures stimulates NE activity in neutrophils, but NE activity is significantly reduced after exposure to CCM from DDR1-knockdown clones (Figure 6H). In addition, heat treatment of NCCM harvested from DDR1 intact cancer cell/collagen I/neutrophils results in a reduction of cancer cell invasion (Figure 6I), further suggesting that a secreted protein is responsible for the observed effect.

**Figure 6.**
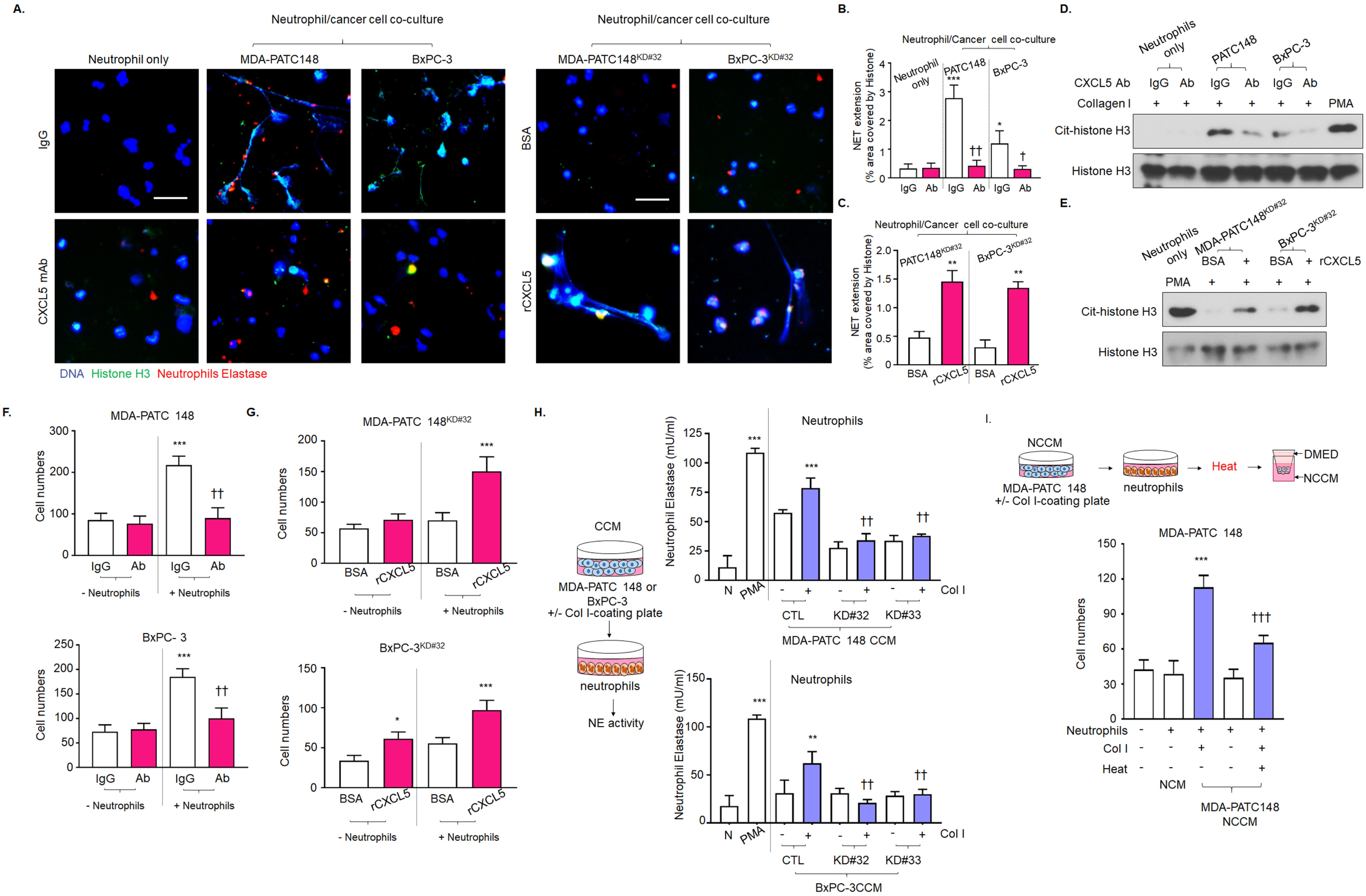
CXCL5 involved in DDR1 mediated NET formation and cancer cell invasion. **A-F** Human neutrophils were co-culture with DDR1-knockdown or re-expression of MDA-PATC 148 or BxPC-3 by matrigel transwell chamber, with or without anti-CXCL5 neutralized antibody or recombinant CXCL5 treatment, for 18 hours. **A** NET structures were analyzed by immunofluorescence staining using DAPI *(blue)*, anti-NE (*red)* and anti-histone *(green)* mAbs. *Scale bar*, 50 μm. **B and C** The NET quantification is displayed as NET histone area (μm^2^) /per filed. **D and E** Cit-histone H3 expression were analyzed by western blotting. **F and G** The number of invaded cells were analyzed by immunofluorescence staining using DAPI and calculated based on the number of cells found in six fields/per chamber. **H** Neutrophils Elastase activity were showed in human neutrophils with CCM treatment for 18 hours. **G** The number of invaded cells were showed in MDA-PATC 148 cells with heat-treated NCCM treatment for 18 hours.

To examine the DDR1-CXCL5 axis and NET formation in human PDAC, we performed immunohistochemistry and immunofluorescence staining on primary human PDAC samples. We found that the areas of high DDR1 expression were associated with high intra-tumoral levels of CXCL5 (black arrows) (Figure 7). Conversely, within the same tumor we observed that regions of low DDR1 expression were associated with decreased CXCL5 expression (red arrow). In addition, we found that the NET-like structures were located around tumor cells (green and red color). Statistically, Pearson *r* correlation analysis indicated a positive association between DDR1 and CXCL5 level in 3 cases of pancreatic patient tumor specimens (case 1: *r* = 0.4423; case 2: *r* = 0.4467; case 3: *r* = 0.4166)

**Figure 7.**
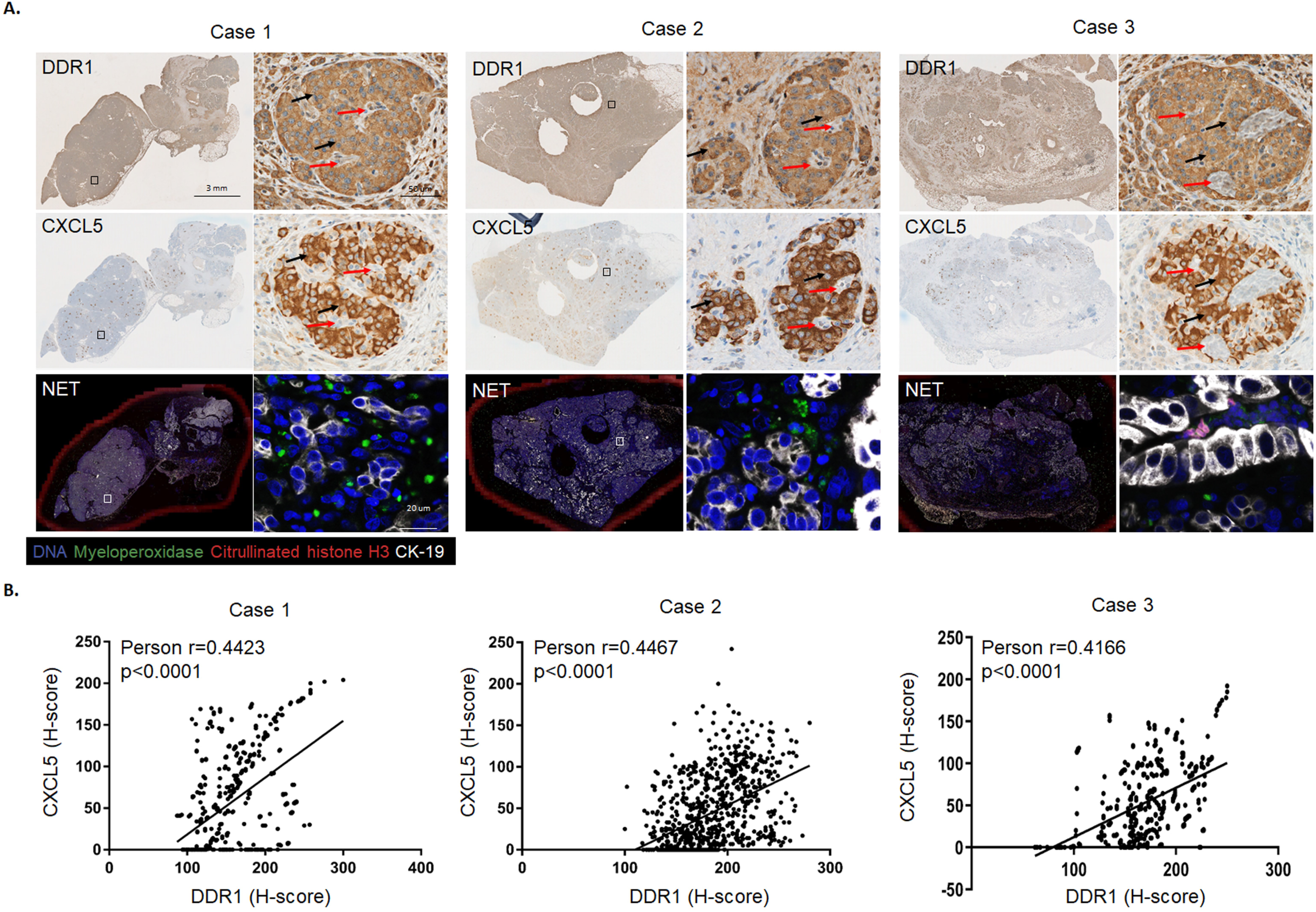
The correlation of DDR1, CXCL5 and NET-like structure in PADC patient samples. **A** *upper and middle panel:* immunohistochemical staining showed DDR1, CXCL5 expression in PDAC patient samples and identified using PE Vectra3. Scale bar, 3mm and 50 μm. *bottom panel:* NET-like structures were analyzed by immunofluorescence staining using DAPI *(blue)*, anti-CK19 (white), anti-MPO *(green)*, anti-citrullinated histone H3 (*red)* mAbs in PDAC patient samples. *Scale bar*, 20 μm. **B** The Pearson correlation showed relationship of DDR1 and CXCL5 by using H-score which quantified the DBA signals by inform software.

### Cancer cells induce CXCL5 and cause NET formation through the DDR1-PKCθ-NF-κB signaling pathway

It has been reported that CXCL5 is a downstream product of the NF-κB pathway ^29^ and that the NF-κB pathway can be activated by DDR1^30^. Therefore, we sought to determine whether the NF-κB pathway is involved in collagen-induced CXCL5 production. A ChIP assay demonstrated that the exposure of MDA-PATC 148 cells to collagen I induced binding of the NF-κB P65 subunit to the CXCL5 promoter and this binding was reduced significantly after the knockdown of DDR1 (Figure 8A). In addition, the exposure of MDA-PATC 148 and BxPC-3 cells to collagen I for 3 h induced NF-κB P65 translocation into the nucleus (Figure 8B and Supplementary Figure 6). Consistent with this, the activation of NF-κB P65 was prevented by genetic DDR1 knockdown in MDA-PATC 148 and BxPC-3 cells (Figure 8B and Supplementary Figure 6). In addition, a phospho-NF-κB pathway array performed after collagen I stimulation of MDA-PATC 148^KD#32^ cells demonstrated the phosphorylation levels of these five proteins with the greatest decrease (20%-25%) in phosphorylation when compared to parental cells: NF-κB P100/52, SYK, NF-κB P65, ZAP-70, and PKCθ (Supplementary Figure 7). Recent studies have shown that SYK is expressed highly in multiple cancer cell types and that SYK kinase activity induces cancer cell migration and metastasis ^31,32^. Thus, we tested whether DDR1 upregulated CXCL5 through NF-κB, SYK, and PKCθ and found that the knockdown of DDR1 decreased collagen I-stimulated NF-κB, SYK, and PKCθ phosphorylation (Figure 8C). Next, we generated an IκB super-repressor mutant clone to block the activation of NF-κB in MDA-PATC 148 (MDA-PATC 148^Iκ B-MUT^) (Supplementary Figure 8) and found that this significantly inhibited collagen I-induced CXCL5 production (Figure 8D). Additionally, pretreatment with SYK and PKC inhibitors prevented collagen I-induced CXCL5 production and inhibited NF-κB P65 activation in MDA-PATC 148 and BxPC-3 cells (Figure 8E and F). In addition, PKC inhibitors blocked the effect of collagen I-induced SYK phosphorylation; in contrast, the SYK inhibitor had no effect on collagen I-induced PKCθ activation (Figure 8F). These results suggest that collagen I-induced CXCL5 production is mediated by a DDR1-PKCθ-SYK-NFκB signaling cascade.

**Figure 8.**
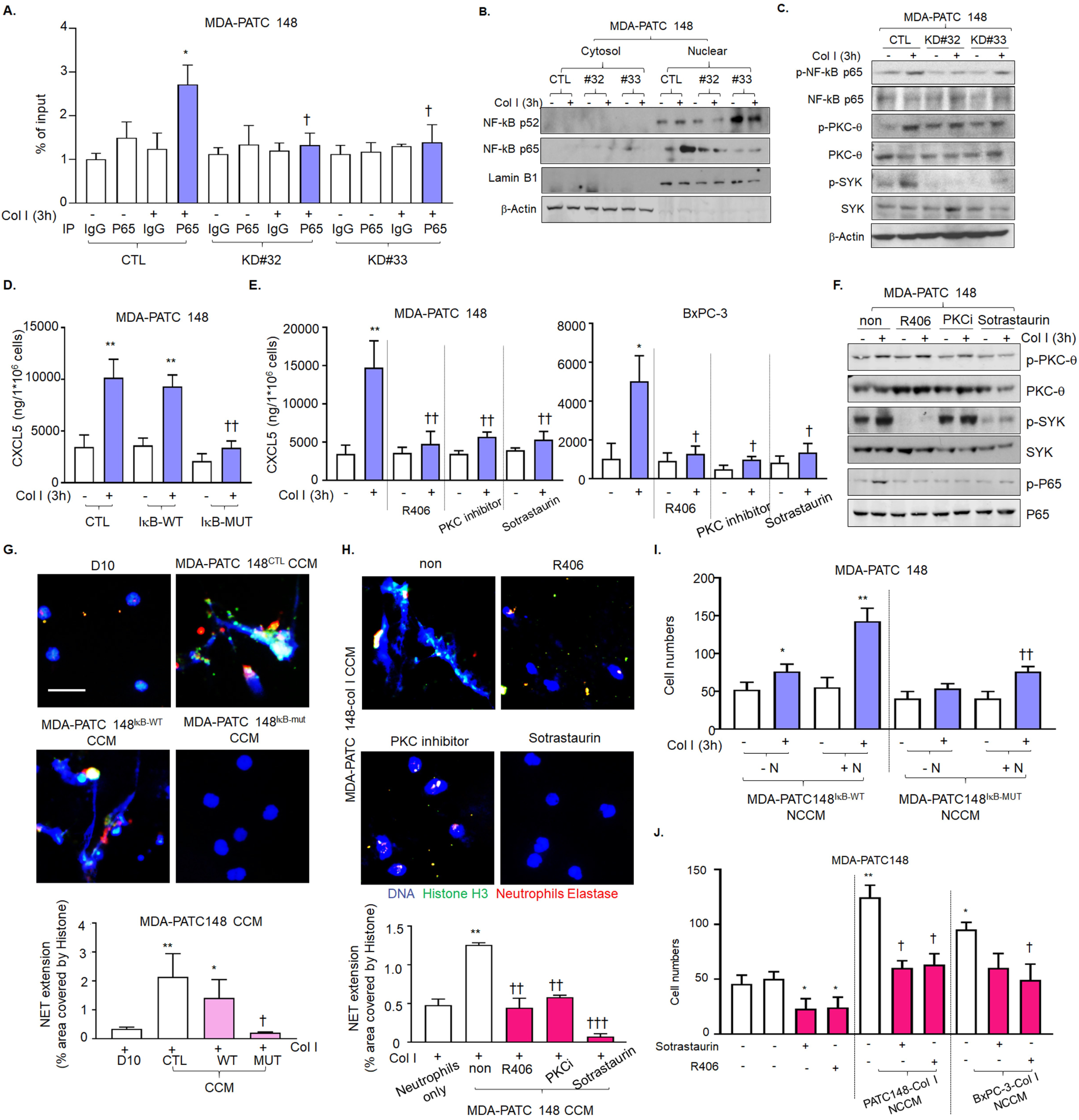
PKCq-SYK-NFkB pathway involved in DDR1 induced CXCL5 production, NET formation form neutrophils and enhanced cancer cell invasion. **A** qPCR results were used to quantify enrichment of NF-kB P65 at the CXCL5 promoter using chromatin immunoprecipitation (ChIP) assay in MDA-PATC 148 cells with DDR1 knockdown. **B** activated NF-kB were analyzed by western blotting using nuclear fraction in MDA-PATC 148 cells with DDR1 knockdown. **C** Phospho-NF-kB P65, phospho-PKCq and phospho-SYK were analyzed by western blotting in MDA-PATC 148 cells with DDR1 knockdown. **D and E** CXCL5 level were analyzed by ELISA **D** in MDA-PATC 148 cells with IkB superrepressor mutation. **E** in MDA-PATC 148 and BxPC-3 cells with or without SYK inhibitor and PKC inhibitor pretreatment. **F** Phospho-NF-kB P65, phospho-PKCq and phospho-SYK were analyzed by western blotting in MDA-PATC 148 cells with or without SYK inhibitor and PKC inhibitor pretreatment. **G and H** NET structures were analyzed by immunofluorescence staining using DAPI *(blue)*, anti-NE (*red)* and anti-histone *(green)* mAbs **G** in MDA-PATC 148 cells with IkB superrepressor mutation. **H** in MDA-PATC 148 cells with or without SYK inhibitor and PKC inhibitor pretreatment.. *Scale bar*, 50 μm. The NET quantification is displayed as NET histone area (μm^2^) /per filed. **I and J** The number of invaded cells were analyzed by immunofluorescence staining using DAPI and calculated based on the number of cells found in six fields /per chamber. I in MDA-PATC 148 cells with NCCM from MDA-PATC 148 cells with IkB superrepressor mutation/neutrophils/collagen I, treatment for 18 hours. J in MDA-PATC 148 cells with NCCM from MDA-PATC 148/ collagenl/SYK or PKC inhibitor, treatment for 18 hours.

To confirm the contribution of the PKCθ-SYK-NFκB axis to DDR1-mediated NET formation, we harvested CCM from MDA-PATC 148^IκB-WT^, MDA-PATC 148^IκB-MUT^ cells and MDA-PATC 148 cells pretreated with SYK or PKC inhibitors after exposure to collagen I for 3 h and incubated neutrophils with the CCM for 18 h. CCM from cancer cells incapable of NF-κB activation was associated with decreased NET formation after collagen I stimulation (Figure 8G). In addition, pretreatment with SYK and PKC inhibitors protected against MDA-PATC 148 CCM-induced NET formation (Figure 8H). Finally, we tested whether DDR1-mediated NET-induced cancer cell invasion was through the PKCθ-SYK-NFκB axis. NCCM harvested from MDA-PATC 148^IκB-WT^/collagen I neutrophil cultures significantly induced cancer cell invasion while NCCM from MDA-PATC 148^IκB-MUT^/collagen I neutrophil cultures was associated with diminished cancer cell invasion (Figure 8I). NCCM from MDA-PATC 148 and BxPC-3 cells pretreated with SYK or PKC inhibitors/collagen I neutrophil cultures also reduced cancer cell invasion (Figure 8J).

11 Together, these results provide evidence that DDR1 stimulates CXCL5 production through a PKCθ-SYK-NFκB signaling cascade and that CXCL5 from cancer cells induces neutrophils to form NETs and thereby enhances metastasis.

### Targeting DDR1 reduces neutrophil-mediated cancer cell invasion

Our previous study demonstrated that the specific DDR1 inhibitor 7rh benzamide can improve the efficacy of standard-of-care chemotherapy in PDAC ^12^. In the current study, we wanted to determine whether inhibition of DDR1 signaling by 7hr benzamide would result in a reduction of CXLC5 and subsequent NET formation. We found that pretreatment with 3 μM of 7rh benzamide in MDA-PATC 148 cells significantly reduced NFκB, PKCθ and SYK phosphorylation as well as the production of CXCL5 (Figure 9A and 9B). As predicted, NET formation and cell invasion was significantly decreased in 7rh benzamide-treated cells (Figure 9C-9F).

**Figure 9.**
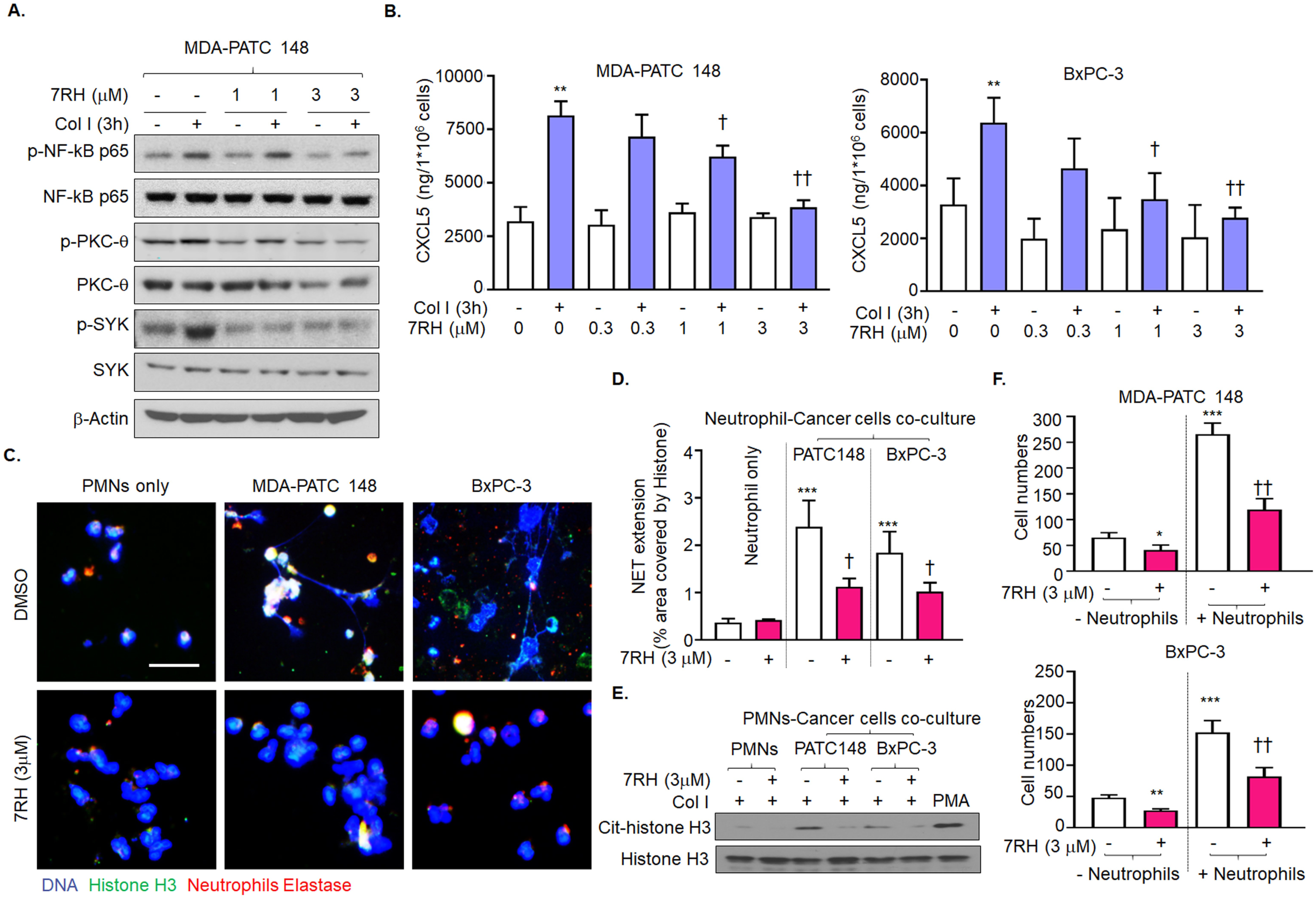
7rh treatment reduced NET formation through inhibition of DDR1-PKCq-SYK-CXCL5 axis and reduced cancer cell invasion. **A and B** MDA-PATC 148 cells and BxPC-3 were pretreated with 7rh for 30 min and then with collagen I for 3 hours. **A** Phospho-NF-kB P65, phospho-PKCq and phospho-SYK were analyzed by western blotting. **B** CXCL5 level were analyzed by ELISA. **C-F** human neutrophils were co-culture with MDA-PATC 148 and BxPC-3 cells by matrigel transwell chamber for 18 hours. **C** NET structures were analyzed by immunofluorescence staining using DAPI *(blue)*, anti-NE *(red)* and anti-histone *(green)* mAbs. *Scale bar*, 50 μm. **D** The NET quantification is displayed as NET histone area (μm^2^) /per filed. **E** Cit-histone H3 expression were analyzed by western blotting. **F** The number of invaded cells analyzed by immunofluorescence staining using DAPI and calculated based on the number of cells found in six fields /per chamber.

## Discussion

Fibrillar collagen is abundant within the microenvironment of primary tumors and is localized adjacent to cancer cells expressing high levels of DDR1^33^. Here we report a novel mechanism by which a collagen receptor, DDR1, on PDAC cells interacts with type I collagen to attract tumor-associated neutrophils, induce NET formation, and facilitate cancer cell invasion and metastasis. In this study, we mechanistically link DDR1 expression in human PDAC with CXCL5 levels, Ly6G^+^ neutrophil infiltration, NET-like structures, and metastatic events. Importantly, in animal models of PDAC, tumors derived from cancer cells lacking DDR1 had fewer NET-like structures and the animals experienced fewer liver metastases, suggesting that cancer cell-derived DDR1 contributes significantly to NET-induced tumor metastasis.

NETs have been reported to link neutrophils and metastasis^21^. Not only neutrophils but also other types of leukocytes, including macrophages, mast cells, and eosinophils, form extracellular traps (a process called ETosis)^34^, but NET formation is the most prominent ETosis in cancer. Cancer cell-mediated cytokines and chemokines can prime neutrophils for NET formation by inducing NADPH oxidase activation to support tumor metastasis. NADPH oxidase and PDA4 inhibitors have been reported to significantly decrease tumor cell invasion, suggesting that NET-mediated tumor cell invasion requires neutrophil NADPH oxidase and PDA4 activity ^21^. Similar to Park *et. al*., we also found that PDA4 and NE activity are important for pancreatic cancer cell-induced NET formation and NET-mediated cancer cell invasion; however, we found that NADPH inhibition had no effect on NET-mediated tumor cell invasion, suggesting that PDAC cells induce NET formation through an NADPH oxidase-independent pathway (Figures 5E, F, and G). It has been shown that CXCR2-related chemokines induce NET formation through the NADPH oxidase-independent pathway ^35^. This is consistent with our findings that CXCL5, a CXCR2 ligand, is expressed by tumor cells in a DDR1-dependent manner and drives neutrophil activation and NET formation (Figures 2 and 6A). Recent studies have suggested that NET-associated DNA meshes can catch circulating tumor cells and enhance cell migration and spread ^19,36^. In addition, NET-associate proteases, NE and MMP9, awaken dormant cancer cells and facilitate cancer cell spreading to different tissues^20,21^. In our studies, decrease in NE activity through DDR1 knockdown, NE inhibitors, or heat treatment all prevented NET-mediated cancer cell invasion, highlighting the contribution of NE in PDAC cell invasion (Figures 5G, 6H, and 6I). Taken together, these data support that collagen I-DDR1 interaction induces CXCL5 production by cancer cells, which promotes neutrophil infiltration and NET formation to drive NET-associated cancer cell migration.

In patients, an elevated serum CXCL5 level has been statistically associated with liver metastasis and poor survival^37,38^. CXCL5 recruits neutrophils into the TME ^25^, and the CXCL5/CXCR2 axis contributes to tumor growth and metastasis through the activation of PI3K/Akt/GSK-3β/Snail signaling to promote EMT^39^. However, little is known about the mechanism of CXCL5 production. Our studies demonstrate that the activation of DDR1 signaling by collagen I induced CXCL5 mRNA and protein expression in PDAC cells *in vitro* and *in vivo*. The STAT3 and NF-κB pathways are involved in DDR1 downstream signaling and have been reported to induce CXCL5 production^29,40,41^. However, in our system, the inhibition of STAT3 by pretreated cucurbitacin I had no effect on collagen I-induced CXCL5 production (Supplementary Figure 9). For this reason, we focused on the NF-κB pathway and identified a putative signaling cascade from DDR1 to CXCL5 expression via the PKCθ-SYK-NFκB axis (Figure 8B-F). This is consistent with descriptions of how NFκB is associated with the hallmarks of cancer, including cancer cell proliferation, protection against apoptosis, and metastasis^42,43^. In addition to NFκB, PKCθ also promotes cancer metastasis by upregulating EMT and the matrix metalloprotease, MMP-1^44,45^, characteristics that are also consistent with our observations.

Constitutive KRAS and NF-κB activation are signature alterations in PDAC^46^. Kras^G12D^-induced cytokines and CXCR2-related chemokines have been reported to facilitate myeloid cell infiltration and tumor progression^47-49^. In addition, we previously showed that Kras^G12D^*-*activated IL-1α/NF-κB /IL-1α and p62 feedforward loops are necessary for the induction and maintenance of NF-κB activity ^50^. Here we found that BxPC-3 cells, which are Kras WT, could also produce CXCL5 to drive NET formation. BxPC-3 cells have been reported to bind with high affinity to ECM proteins, including collagen I and possess invasive capabilities consistent with our observations ^51^. Nevertheless, it is possible that Kras-mediated NF-κB could drive CXCL5 production, neutrophil infiltration, and NET formation. MDA-PATC 148 cells, which are Kras-mutant, generated higher levels of CXCL5 with associated NET formation and invasion when compared with BxPC-3 cells (Figures 2C, 5A-D). Although Kras^G12D^ may intrinsically drive NF-κB–CXCL5-mediated NET, our work has identified a novel mechanism by which an abundant stromal molecule, collagen I, can interact with cancer cells through a targetable receptor (DDR1) to induce CXCL5 production, NET formation, and NET-mediated cancer cell invasion even in the absence of a Kras mutation.

Therapeutic strategies must also account for stromal cells in the TME, which also play an important role in the promotion of metastasis ^52^. In the current study, we focus on DDR1 and two stromal components: collagen I and tumor-associated neutrophils. DDR1 amplification is commonly observed in various cancers ^53^. In addition, the DDR1 signaling pathway is significantly deregulated in aggressive cancers ^54^. We observed higher levels of DDR1 in metastatic cell lines compared with matched primary cell lines (Figure 1A). Emerging evidence suggests a crucial role for DDR1 in the progression and metastasis of various solid tumors; as such, targeting DDR1 represents a promising therapeutic approach ^55-57^. This is consistent with our repeated observations of a reduction of cell invasion and metastasis after pharmacologic or genetic inhibition of DDR1 signaling (Figures 1C-F, 5D, 9, and Supplementary Figure 9). The role of neutrophils in solid tumors remains, however, poorly defined. Although tumor-associated neutrophils support cancer cell progression and invasion, they can also kill cancer cells and bacteria within the TME ^58^. For example, the American Society of Clinical Oncology recommends that patients who are undergoing certain chemotherapy regimens receive prophylactic treatment with granulocyte colony-stimulating factor, which stimulates neutrophil production to counteract neutropenia^59^. It is possible, however, that inducing neutrophil production may increase the risk of metastatic spread. Recent work suggests that NET production might awaken clinically dormant cancer cells after chemotherapy and drive them to become metastatic^20^. Our previous study suggested that a small-molecule inhibitor of DDR1 signaling, 7rh benzamide, would improve the efficacy of standard-of-care chemotherapy in patients with PDAC^12^. In this study we found that 7rh significantly reduced cancer cell-induced NET formation in support of our prior observations. Taken together, we suggest that 7rh, when given in therapeutic doses, could increase the sensitivity of cancer cells to conventional chemotherapy and inhibit liver metastasis by blocking NET formation. DDR1 is critical in this process and might also present a valuable pharmacological target in the treatment of patients with PDAC.

## Materials and Methods

### Cell lines

Pancreatic cancer cell lines—MDA-PATC 43, 50, 53, 66, 69, 102, 108, 124, 148, 153, and 216— were generated in our laboratory from PDX tumors^60^. The MDA-PATC 148LM and MDA-PATC 153LM cell lines were generated from liver metastases of MDA-PATC 148 and 153 cells *orthotopically implanted* into nude mice respectively. PANC-1 and BxPC-3 cell lines were obtained from the American Type Culture Collection (Manassas, VA). WM8865 mouse cells were generated from *Kras*^*LSL*.*G12D/+*^; *p53*^*R172H/+*^; *Pdx1*^*CreTg/+*^ (*KP*^*wm*^*C*) mice in our laboratory. All cells were cultured in DMEM that contained 10% FBS, penicillin (100 units/mL), and streptomycin (100 µg/mL; D10 medium), and had been tested monthly for mycoplasma and found negative.

### Animal studies

All animal experiments were conducted according to the National Institutes of Health’s Guide for the Care and Use of Laboratory Animals. The study protocol was approved by the University of Texas MD Anderson Cancer Center’s institutional animal care and use committee.

We randomly assigned 8-week-old nude mice and C57BL/6J mice from the Jackson Laboratory (Bar Harbor, ME) to different groups: nude mice with MDA-PATC 148^CTL^ and MDA-PATC 148^KD#32^; and C57BL/6J mice with KP^wm^C^CTL^, KP^wm^C^KD#588^ and KP^wm^C^KD#809^. We resuspended 1 × 10^5^ cells in 25 µL 1X PBS and added one volume of Matrigel (Corning, CB356253). The suspension was then directly injected into the pancreas of mice. Nine weeks after cancer cell injection, all mice were euthanized, and we collected their plasma and pancreatic, liver, and spleen tissues.

### PDX tumors and tissue microarrays

We followed our previously published protocol for the heterotopic engraftment of pancreatic patient tumors into immunodeficient mice^61^. After harvesting the tissues from the euthanized mice, we embedded them in paraffin and used core samples to construct a tissue microarray.

### Isolation of human neutrophils

To isolate human neutrophils, we obtained peripheral venous blood samples (10 mL) that were collected from untreated pancreatic cancer patients and stored at MD Anderson Cancer Center (IRB: PA11-0670). Isolation of the neutrophils was modified as described previously^62^. After the plasma and platelets were removed from the blood, the granulocytes were isolated using a Ficoll-Paque PLUS (GE Healthcare, 17-1440-03) with density-gradient centrifugation, and the erythrocytes were removed by two rounds of hypotonic lysis with a red blood cell lysis buffer. The viability and purity of the neutrophils were determined by anti-human CD66b PE (clone G10F5; BD Biosciences, 561650) and anti-human CD16 FITC (clone 3G8; BD Biosciences, 560996) double staining using FACS (Supplementary Figure 10).

### *In vitro* cell invasion assay

Our cell invasion assay was modified as described previously ^21^. We suspended 1 × 10^5^ neutrophils in 100 µL of D10 medium and seeded the cells on poly-L-lysine–coated coverslips in a 24-well culture plate for 30 min. We then removed the medium and non-adhered neutrophils and added 700 µL of D10 medium with or without IgG, anti-CXCL5 antibody (Abcam, ab9802), BSA, human recombinant CXCL5 (Abcam, ab9803), DNase I (1.5U), apocynin (10 µM; Abcam, ab120615), the PAD4 inhibitor, Cl-amidine (200 µM; Cayman Chemical, 1043444-18-3), and sivelestat (10 µM; ApexBio, B6189). We suspended 2 × 10^5^ cancer cells in 500 µL of serum-free DMEM medium (D0 medium) with or without 7rh, IgG, anti-CXCL5 antibody, BSA, human recombinant CXCL5, DNase I, NADPH oxidase inhibitor, apocynin, PAD4 inhibitor, Cl-amidine, and the neutrophil elastase inhibitor sivelestat and then added the cells to a rehydrated Matrigel chamber (Corning, 354480). After the cells had been cultured for 18 h at 37°C, the invading cancer cells were fixed with 100% cold methanol and stained with 0.05 mg/mL 4,6-diamidino-2-phenylindole (DAPI; BD Biosciences, 564907). The invading cells in each chamber were counted under a fluorescence microscope, and the average number of cells was calculated based on the number of cells found in six fields per chamber.

To generate cancer cell conditioned medium (CCM), we suspended 2 × 10^6^ cancer cells cultured in 2 mL of D0 medium with or without the SYK inhibitor R406 (10 µM; BioVision, 9682); the PKC inhibitor (30 µM; Santa Cruz Biotechnology, sc-3007); another PKC inhibitor, sotrastaurin (2 nM; Selleck Chemicals, S2791); and different dosages of the DDR1 kinase inhibitor, 7rh, for 30 min at 37°C. We then seeded the cells on collagen I-coated 6-well culture plate for 3 h. Next, we removed and stored the medium at -80°C. Neutrophil conditioned medium (NCM) was prepared from 1 × 10^6^ neutrophils cultured in 1 mL of D10 medium with or without phorbol 12-myristate 13-acetate (PMA; Sigma-Aldrich, now MilliporeSigma, P8139) for 4 h; human recombinant CXCL5 for 18 h; then we collected the and stored the medium at -80°C. Neutrophils exposed to cancer cell conditioned media (NCCM) was prepared from 1 × 10^6^ neutrophils cultured in CCM for 18 h; then we collected the medium with or without heat denaturing (95°C) for 5 min, and stored it at -80°C. The NCM or NCCM was added to a 24-well culture plate, and the cancer cells were seeded in D0 medium in a rehydrated Matrigel chamber. After the cells had been cultured for 18 h at 37°C, the invading cancer cells were fixed with 100% cold methanol and stained with 0.05 mg/mL DAPI. The invading cells in each chamber were counted under a fluorescence microscope, and the average number of cells was calculated based on the number of cells found in six fields per chamber.

### Detection of NET structure by immunofluorescence staining

Neutrophils grown on coverslips were fixed, permeabilized with 4% paraformadehyde and 0.5% Triton X-100 in PBS, and blocked in PBS containing 2.5% FBS and 2.5% BSA for 1 h at room temperature. The neutrophils were then incubated overnight at 4°C with primary antibodies: anti-histone H3 (1:50 dilution; Cell Signaling Technology, 3680) and either anti-NE (1:100 dilution; Abcam, ab68672) or anti-myeloperoxidase (1:150 dilution; R&D Systems). After being washed with PBS, secondary antibodies conjugated with Alex488 or TRITC were applied to detect primary Ab (1:250 dilution), and DAPI was used as a counterstain. NET images, defined as areas of co-localized DNA, myeloperoxidase, and citrullinated histone H3, were observed in five fields of each coverslip using a fluorescence microscope, and histone area was used to quantify NET extension using ImageJ software.

For tissue histology, paraffin-embedded sections of tumors were deparaffinized and rehydrated; antigen retrieval was in citrate buffer (10 mM sodium citrate, 0.05% Tween 20, pH 6.0), and the sections were blocked in PBS containing 3% BSA for 1 h. Then the tissue sections were incubated overnight at 4°C with antibodies against myeloperoxidase (1:150 dilution; R&D Systems, AF3667), citrullinated histone H3 antibody (1:100 dilution; Abcam, ab5103), and CK-19 (1:500; Abcam). The tissue sections were then stained with secondary antibodies conjugated with Alex488; TRITC or Cy5 to detect primary Ab (1:250 dilution), and DAPI was used as a counterstain. NET structure was identified using an automated multispectral imaging microscope (Vectra3, PerkinElmer) and analyzed by using inForm software at the MD Anderson Cancer Center’s Flow Cytometry and Cellular Imaging Core Facility.

### Immunohistochemistry

Paraffin-embedded sections of tissue obtained from PDX tumors, human tumors in tissue microarrays, or tumors derived from MDA-PATC 148 cells with knockdown DDR1 tumors were deparaffinized and rehydrated; antigen retrieval was in citrate buffer (10 mM sodium citrate, 0.05% Tween 20, pH 6.0). Sections were treated with 3% H_2_O_2_, blocked with Fc Receptor blocker (Innovex), and incubated with 1x blocking buffer (5% BSA in PBS) for 1 h. Then the tissue sections were incubated overnight at 4°C with antibodies against human DDR1 (R&D Systems, AF2396), CXCL5 (Abcam, ab9802), Ly6G (BD Pharmingen, 551459) antibodies. Biotinylated secondary antibodies (VECTASTAIN ABC kit, Vector Labs) were used for primary antibody detection following the manufacturer’s protocols. Sections were counterstained with hematoxylin. Images were identified using an automated multispectral imaging microscope (Vectra3) and analyzed by using inForm software at the MD Anderson Cancer Center’s Flow Cytometry and Cellular Imaging Core Facility (Supplementary Figure 11).

### Short hairpin RNAs

Vectors expressing short hairpin RNA (shRNA) against human DDR1 (#32: 5’-GACAGCCCATCACCTCTAA-3’; #33: 5’-CAGGTCCACTGTAACAACA-3’) and mouse DDR1 (#588: 5’-TGCAGCTAGAACTTCGCAA-3’; #809: 5’-AGGTCCTTGGTTACTCTTC-3’) were generated using the pGIPZ-shRNA plasmids and were packaged into lentiviral particles at the MD Anderson Cancer Center’s shRNA and ORFeome Core. Viruses were transfected into BxPC-3 and MDA-PATC 148 cell lines. Puromycin (2 μg/mL) was used to remove nontransfected cells.

### DDR1 overexpression plasmids

We packaged pLX304-Blast-V5 vector-expressing human DDR1 ORFs (Dharmacon) into lentiviral particles at MD Anderson Cancer Center’s shRNA and ORFeome Core. Viruses were transfected into Panc1, Miacapa-2, and MDA-PATC 53, 66, 148#32, and 153 cell lines. Blasticidin (0.3 µg/ml) was used to remove nontransfected cells.

### IκB super-repressor mutant plasmids

IκB super-repressor mutant plasmids were generated using pBABE vectors and were packaged into retrovirus particles at MD Anderson Cancer Center’s shRNA and ORFeome Core. Viruses were transfected into MDA-PATC 148 cells. Puromycin (2 μg/mL) was used to remove nontransfected cells.

### Chemokine and NF-κB phospho antibody arrays

For the chemokine array, we cultured 1 × 10^6^ cancer cells in 1 mL of D0 medium for 16 h at 37°C. Then we collected the supernatants and cell lysates so that we could detect the chemokines using a Proteome Profiler Human Chemokine Array Kit (R&D Systems, ARY017).

For the NFκB phospho antibody array, we suspended 2 × 10^6^ cancer cells in 2 mL of D0 medium and seeded the cells on collagen I-coated 6-well culture plates for 3 h. The cell lysates were then collected, and we detected the NFκB pathway using an NFκB Phospho Antibody Array (Full Moon BioSystems, KAS02).

### ELISA

We suspended 2 × 10^5^ cancer cells in 200 µL of D0 medium and pretreated the cells with or without SYK inhibitor R406 and the PKC inhibitor sotrastaurin for 30 min at 37°C. Then cells were seeded on control or collagen I-coated 96-well plates for 3 h. Next, the culture supernatants were collected and stored at -80°C. Cells and tumor tissues were lysed and quantified with a Bio-Rad protein assay kit. Culture supernatants, cell lysates, tumor lysates, and plasma were assayed for CXCL5 using a human/mouse CXCL5 immunoassay kit (BioLegend, 440904).

### Real-time PCR

We quantified mRNA expression using real-time PCR. RNA was prepared using TRIzol Reagent (Invitrogen). We synthesized cDNA using the StepOne Real-Time PCR System (Bio-Rad, 1708840) and analyzed it using StepOne Software v2.2.1 (Bio-Rad). Each sample was tested in triplicate for CXCL1, CXCL2, CXCL3, CXCL5, CXCL6, CXCL7, and CXCL8, and results were normalized by real-time PCR of the cDNA with glyceraldehyde 3-phosphate dehydrogenase (GAPDH). The primers are shown in Table S1.

### Detection of neutrophil infiltration in tumors using FACS

Mice were perfused with a PBS buffer containing heparin (10 U/mL) prior to being euthanized. Then 100 mg of tumor was collected and isolated in single cells by the buffer containing 5 mg/ml collagenase type IV (Gibco, 17104-019) and 1 mg/ml dispase II (Gibco, 17105-041) at 37 °C for 2 h. After 2 h, tumor homogenates were passed through a 100 µm cell strainer and lysed with a red blood cell lysis buffer to remove the remaining red blood cells. The homogenates were then resuspended in PBS buffer and stained with anti-mouse CD45.1 antibody conjugated with Brilliant Violet 421 (clone A20; BioLegend, 103134), CD11b monoclonal antibody conjugated with APC (M1/70; eBioscience, 17-0112-82), and anti-mouse Ly6G antibody conjugated with APC/Cy7 (clone1A8; BioLegend, 127624). Stained cells were washed twice with FACS buffer and were resuspended in PBS buffer and analyzed. CD45^+^CD11b^+^Ly6G^+^ neutrophil infiltration in tumors was identified using a Beckman Coulter Gallios flow cytometer (Gallios 561) at the MD Anderson Cancer Center’s Flow Cytometry and Cellular Imaging Core Facility. All data were processed using FlowJo software, version 10 (Treestar).

### Nuclear protein extraction and immunoblotting

Cytosolic and nuclear fractions were modified as described previously^63^. Cell pellets were lysed in buffer A (20 mM HEPES at pH 7, 10 mM KCl, 2 mM MgCl2, 0.5% NP-40, 1 mM NaF, 1 mM Na_3_VO_4_, 1 mM phenylmethylsulfonyl fluoride, and 1 μg/mL aprotinin), homogenized in a glass dounce, and centrifuged at 1500 x *g* for 10 min. The supernatant was the cytosolic fraction. The nuclear pellet was isolated and washed three times in buffer A. The nuclei were sonicated in RIPA buffer and centrifuged at 15,000 x *g* for 20 min. Proteins were quantified with a Bio-Rad protein assay kit. Equal amounts of protein from each sample were subjected to SDS PAGE and transferred to a polyvinylidene membrane (Invitrogen). The membrane was blocked with 5% skim milk in tris-buffered saline with 0.1% Tween 20 for 1 h. It was then hybridized with DDR1 (D1G6; Cell Signaling, 5583), SYK (Cell Signaling, 2712), phospho-SYK (Cell Signaling, 2710), PKCθ (Cell Signaling, 13643), phospho-PKCθ (Cell Signaling, 9376), NF-κB p65 (Cell Signaling, 6956), phospho-NF-κB p65 (Cell Signaling, 3033), NF-κB p50/100, and β-actin (Sigma-Aldrich, A5316) primary antibodies. The membrane was then incubated with horseradish peroxidase–conjugated secondary antibodies, and the bands were visualized using Western Lightning Plus-ECL (PerkinElmer).

### Chromatin immunoprecipitation assay

Overall, chromatin immunoprecipitation (ChIP) assays were modified as described previously^64^. An antibody against activated NF-κB p65 antibody (Abcam, ab19870) was used for the ChIP assay. Binding of active p65 to the CXCL5 promoter was quantified by real-time PCR in MDA-PATC 148 cells, with or without DDR1 knockdown. The primers are shown in Table S1.

### Statistical analysis

For *in vitro* cell invasion assay, NET extension, and cell stimulation assay, an unpaired *t* test was used to compare invaded cell number, histone H3 citrullination, and cytokine production between groups. Fisher’s exact test was used to evaluate the number of mice with liver metastasis.

*In vivo* tissue histology scores were compared by the Mann-Whitney *U* test (two-tailed). A Pearson *r* correlation was used to evaluate the association of DDR1/CXCL5, DDR1/Ly6G, and CXCL5/Ly6G histology scores. These analyses were performed using GraphPad Prism version 8.

### Study approval

This study was approved by the institutional review board and the institutional animal care and use committee of the University of Texas MD Anderson Cancer Center. Patients gave written informed consent for participation. All animal experiments were conducted according to the National Institutes of Health’s Guide for the Care and Use of Laboratory Animals.

## Author contributions

**Conception and design:** J.B. Fleming, J. Deng

**Development of methodology**:J.B. Fleming, J. Deng, Y. Kang, C-C. Cheng

**Acquisition of data (provided animals, acquired and managed patients, provided facilities, etc**.):J.B. Fleming, J. Deng, Y. Kang, X. Li

**Analysis and interpretation of data (e**.**g**., **statistical analysis, biostatistics, computational analysis):**J. Deng, Y. Kang, X. Li, B. Dai

**Writing, review, and/or revision of the manuscript**: J.B. Fleming, J. Deng, H. Huang, R.A. Brekken

**Administrative, technical, or material support (i**.**e**., **reporting or organizing data, constructing databases):** J.B. Fleming, J. Deng, C-C. Cheng, B. Dai, M.H. Katz, T. Men, M.P. Kim, E.A. Koay, R.A. Brekken

**Study supervision**: J.B. Fleming

## Acknowledgments

This work was supported by grants from the NCI U54 CA210181 (JBF, EAK, and RAB) and R01 CA192381 (RAB). This work was also supported by resources of the MD Anderson Flow Cytometry and Cellular Imaging Core Facility and Functional Genomics Core under NIH/NCI awards (P30CA076292, H. Lee Moffitt Cancer Centers; and P30CA016672, MD Anderson). The authors declare no competing financial interests. They would like to thank the education and training center at MD Anderson and UT Southwestern Medical Center for help in editing this article.

## Abbreviations

CCM: cancer cell conditioned medium
ChIP: chromatin immunoprecipitation
NCCM: cancer cell neutrophil conditioned medium
DDR: discoid domain receptor
ECM: extracellular matrix
EMT: epithelial-mesenchymal transition
ENA-78: epithelial-derived neutrophil-activating peptid 78
GAPDH: glyceraldehyde 3-phosphate dehydrogenase
GEMM: genetically engineered mouse model
MPO: myeloperoxidase
NADPH: nicotinamide adenine dinucleotide phosphate
NCCM: neutrophils exposed to cancer cell conditioned media
NCM: neutrophil conditioned medium
NE: neutrophil elastase
NETs: neutrophil extracellular traps
PAD4: peptidylarginine 4
PDAC: pancreatic ductal adenocarcinoma
PDX: patient-derived xenograft
PMA: phorbol myristate acetate
TAMs: tumor-associated macrophages
TANs: tumor-associated neutrophils
TME: tumor microenvironment

## Notes

### Competing Interest Statement

The authors have declared no competing interest.

## References

1 Neoptolemos, J. P. et al. Therapeutic developments in pancreatic cancer: current and future perspectives. Nat Rev Gastroenterol Hepatol 15, 333–348, doi:10.1038/s41575-018-0005-x (2018).

2 Rahib, L. et al. Projecting cancer incidence and deaths to 2030: the unexpected burden of thyroid, liver, and pancreas cancers in the United States. Cancer Res 74, 2913–2921, doi:10.1158/0008-5472.CAN-14-0155 (2014).

3 Hosein, A. N., Brekken, R. A. & Maitra, A. Pancreatic cancer stroma: an update on therapeutic targeting strategies. Nat Rev Gastroenterol Hepatol, doi:10.1038/s41575-020-0300-1 (2020).

4 Huang, H., Du, W. & Brekken, R. A. Extracellular Matrix Induction of Intracellular Reactive Oxygen Species. Antioxid Redox Signal 27, 774–784, doi:10.1089/ars.2017.7305 (2017).

5 Huang, H., Wright, S., Zhang, J. & Brekken, R. A. Getting a grip on adhesion: Cadherin switching and collagen signaling. Biochim Biophys Acta Mol Cell Res 1866, 118472, doi:10.1016/j.bbamcr.2019.04.002 (2019).

6 Leitinger, B. Discoidin domain receptor functions in physiological and pathological conditions. Int Rev Cell Mol Biol 310, 39–87, doi:10.1016/B978-0-12-800180-6.00002-5 (2014).

7 Valiathan, R. R., Marco, M., Leitinger, B., Kleer, C. G. & Fridman, R. Discoidin domain receptor tyrosine kinases: new players in cancer progression. Cancer Metastasis Rev 31, 295–321, doi:10.1007/s10555-012-9346-z (2012).

8 Huo, Y. et al. High expression of DDR1 is associated with the poor prognosis in Chinese patients with pancreatic ductal adenocarcinoma. J Exp Clin Cancer Res 34, 88, doi:10.1186/s13046-015-0202-1 (2015).

9 Shintani, Y. et al. Collagen I-mediated up-regulation of N-cadherin requires cooperative signals from integrins and discoidin domain receptor 1. J Cell Biol 180, 1277–1289, doi:10.1083/jcb.200708137 (2008).

10 Huang, H. et al. Up-regulation of N-cadherin by Collagen I-activated Discoidin Domain Receptor 1 in Pancreatic Cancer Requires the Adaptor Molecule Shc1. J Biol Chem 291, 23208–23223, doi:10.1074/jbc.M116.740605 (2016).

11 Ruggeri, J. M. et al. Discoidin Domain Receptor 1 (DDR1) Is Necessary for Tissue Homeostasis in Pancreatic Injury and Pathogenesis of Pancreatic Ductal Adenocarcinoma. Am J Pathol, doi:10.1016/j.ajpath.2020.03.020 (2020).

12 Aguilera, K. Y. et al. Inhibition of Discoidin Domain Receptor 1 Reduces Collagen-mediated Tumorigenicity in Pancreatic Ductal Adenocarcinoma. Mol Cancer Ther 16, 2473–2485, doi:10.1158/1535-7163.MCT-16-0834 (2017).

13 Lambert, A. W., Pattabiraman, D. R. & Weinberg, R. A. Emerging Biological Principles of Metastasis. Cell 168, 670–691, doi:10.1016/j.cell.2016.11.037 (2017).

14 Kim, J. & Bae, J. S. Tumor-Associated Macrophages and Neutrophils in Tumor Microenvironment. Mediators Inflamm 2016, 6058147, doi:10.1155/2016/6058147 (2016).

15 Kolaczkowska, E. & Kubes, P. Neutrophil recruitment and function in health and inflammation. Nat Rev Immunol 13, 159–175, doi:10.1038/nri3399 (2013).

16 Papayannopoulos, V. Neutrophil extracellular traps in immunity and disease. Nat Rev Immunol 18, 134–147, doi:10.1038/nri.2017.105 (2018).

17 Cedervall, J. & Olsson, A. K. Immunity Gone Astray - NETs in Cancer. Trends Cancer 2, 633–634, doi:10.1016/j.trecan.2016.10.012 (2016).

18 Tohme, S. et al. Neutrophil Extracellular Traps Promote the Development and Progression of Liver Metastases after Surgical Stress. Cancer Res 76, 1367–1380, doi:10.1158/0008-5472.CAN-15-1591 (2016).

19 Cools-Lartigue, J. et al. Neutrophil extracellular traps sequester circulating tumor cells and promote metastasis. J Clin Invest, doi:10.1172/JCI67484 (2013).

20 Albrengues, J. et al. Neutrophil extracellular traps produced during inflammation awaken dormant cancer cells in mice. Science 361, doi:10.1126/science.aao4227 (2018).

21 Park, J. et al. Cancer cells induce metastasis-supporting neutrophil extracellular DNA traps. Sci Transl Med 8, 361ra138, doi:10.1126/scitranslmed.aag1711 (2016).

22 Yang, L. et al. DNA of neutrophil extracellular traps promotes cancer metastasis via CCDC25. Nature 583, 133–138, doi:10.1038/s41586-020-2394-6 (2020).

23 Demers, M. & Wagner, D. D. Neutrophil extracellular traps: A new link to cancer-associated thrombosis and potential implications for tumor progression. Oncoimmunology 2, e22946, doi:10.4161/onci.22946 (2013).

24 Sivanandham, R. et al. Neutrophil extracellular trap production contributes to pathogenesis in SIV-infected nonhuman primates. J Clin Invest 128, 5178–5183, doi:10.1172/JCI99420 (2018).

25 Zhou, S. L. et al. Overexpression of CXCL5 mediates neutrophil infiltration and indicates poor prognosis for hepatocellular carcinoma. Hepatology 56, 2242–2254, doi:10.1002/hep.25907 (2012).

26 Papayannopoulos, V., Metzler, K. D., Hakkim, A. & Zychlinsky, A. Neutrophil elastase and myeloperoxidase regulate the formation of neutrophil extracellular traps. J Cell Biol 191, 677–691, doi:10.1083/jcb.201006052 (2010).

27 Li, P. et al. PAD4 is essential for antibacterial innate immunity mediated by neutrophil extracellular traps. J Exp Med 207, 1853–1862, doi:10.1084/jem.20100239 (2010).

28 Brinkmann, V. et al. Neutrophil extracellular traps kill bacteria. Science 303, 1532–1535, doi:10.1126/science.1092385 (2004).

29 Roca, H. et al. Apoptosis-induced CXCL5 accelerates inflammation and growth of prostate tumor metastases in bone. J Clin Invest 128, 248–266, doi:10.1172/JCI92466 (2018).

30 Das, S. et al. Discoidin domain receptor 1 receptor tyrosine kinase induces cyclooxygenase-2 and promotes chemoresistance through nuclear factor-kappaB pathway activation. Cancer Res 66, 8123–8130, doi:10.1158/0008-5472.CAN-06-1215 (2006).

31 Luangdilok, S. et al. Syk tyrosine kinase is linked to cell motility and progression in squamous cell carcinomas of the head and neck. Cancer Res 67, 7907–7916, doi:10.1158/0008-5472.CAN-07-0331 (2007).

32 Ghotra, V. P. et al. SYK is a candidate kinase target for the treatment of advanced prostate cancer. Cancer Res 75, 230–240, doi:10.1158/0008-5472.CAN-14-0629 (2015).

33 Yuge, R. et al. Silencing of Discoidin Domain Receptor-1 (DDR1) Concurrently Inhibits Multiple Steps of Metastasis Cascade in Gastric Cancer. Transl Oncol 11, 575–584, doi:10.1016/j.tranon.2018.02.003 (2018).

34 Pertiwi, K. R. et al. Extracellular traps derived from macrophages, mast cells, eosinophils and neutrophils are generated in a time-dependent manner during atherothrombosis. J Pathol 247, 505–512, doi:10.1002/path.5212 (2019).

35 Marcos, V. et al. CXCR2 mediates NADPH oxidase-independent neutrophil extracellular trap formation in cystic fibrosis airway inflammation. Nat Med 16, 1018–1023, doi:10.1038/nm.2209 (2010).

36 Cedervall, J. et al. Neutrophil Extracellular Traps Accumulate in Peripheral Blood Vessels and Compromise Organ Function in Tumor-Bearing Animals. Cancer Res 75, 2653–2662, doi:10.1158/0008-5472.CAN-14-3299 (2015).

37 Li, A. et al. Overexpression of CXCL5 is associated with poor survival in patients with pancreatic cancer. Am J Pathol 178, 1340–1349, doi:10.1016/j.ajpath.2010.11.058 (2011).

38 Kawamura, M. et al. CXCL5, a promoter of cell proliferation, migration and invasion, is a novel serum prognostic marker in patients with colorectal cancer. Eur J Cancer 48, 2244–2251, doi:10.1016/j.ejca.2011.11.032 (2012).

39 Zhou, S. L. et al. CXCR2/CXCL5 axis contributes to epithelial-mesenchymal transition of HCC cells through activating PI3K/Akt/GSK-3beta/Snail signaling. Cancer Lett 358, 124–135, doi:10.1016/j.canlet.2014.11.044 (2015).

40 Seo, M. C. et al. Discoidin domain receptor 1 mediates collagen-induced inflammatory activation of microglia in culture. J Neurosci Res 86, 1087–1095, doi:10.1002/jnr.21552 (2008).

41 Gao, H. et al. Multi-organ Site Metastatic Reactivation Mediated by Non-canonical Discoidin Domain Receptor 1 Signaling. Cell 166, 47–62, doi:10.1016/j.cell.2016.06.009 (2016).

42 Park, M. H. & Hong, J. T. Roles of NF-kappaB in Cancer and Inflammatory Diseases and Their Therapeutic Approaches. Cells 5, doi:10.3390/cells5020015 (2016).

43 Karin, M. NF-kappaB as a critical link between inflammation and cancer. Cold Spring Harb Perspect Biol 1, a000141, doi:10.1101/cshperspect.a000141 (2009).

44 Zafar, A. et al. Chromatinized protein kinase C-theta directly regulates inducible genes in epithelial to mesenchymal transition and breast cancer stem cells. Mol Cell Biol 34, 2961–2980, doi:10.1128/MCB.01693-13 (2014).

45 Sokolova, O., Vieth, M. & Naumann, M. Protein kinase C isozymes regulate matrix metalloproteinase-1 expression and cell invasion in Helicobacter pylori infection. Gut 62, 358–367, doi:10.1136/gutjnl-2012-302103 (2013).

46 Bailey, P. et al. Genomic analyses identify molecular subtypes of pancreatic cancer. Nature 531, 47–52, doi:10.1038/nature16965 (2016).

47 Lesina, M. et al. RelA regulates CXCL1/CXCR2-dependent oncogene-induced senescence in murine Kras-driven pancreatic carcinogenesis. J Clin Invest 126, 2919–2932, doi:10.1172/JCI86477 (2016).

48 Mizukami, Y. et al. Induction of interleukin-8 preserves the angiogenic response in HIF-1alpha-deficient colon cancer cells. Nat Med 11, 992–997, doi:10.1038/nm1294 (2005).

49 Ancrile, B., Lim, K. H. & Counter, C. M. Oncogenic Ras-induced secretion of IL6 is required for tumorigenesis. Genes Dev 21, 1714–1719, doi:10.1101/gad.1549407 (2007).

50 Ling, J. et al. KrasG12D-induced IKK2/beta/NF-kappaB activation by IL-1alpha and p62 feedforward loops is required for development of pancreatic ductal adenocarcinoma. Cancer Cell 21, 105–120, doi:10.1016/j.ccr.2011.12.006 (2012).

51 Deer, E. L. et al. Phenotype and genotype of pancreatic cancer cell lines. Pancreas 39, 425–435, doi:10.1097/MPA.0b013e3181c15963 (2010).

52 Guo, S. & Deng, C. X. Effect of Stromal Cells in Tumor Microenvironment on Metastasis Initiation. Int J Biol Sci 14, 2083–2093, doi:10.7150/ijbs.25720 (2018).

53 Cerami, E. et al. The cBio cancer genomics portal: an open platform for exploring multidimensional cancer genomics data. Cancer Discov 2, 401–404, doi:10.1158/2159-8290.CD-12-0095 (2012).

54 Beltran, H. et al. Divergent clonal evolution of castration-resistant neuroendocrine prostate cancer. Nat Med 22, 298–305, doi:10.1038/nm.4045 (2016).

55 Ambrogio, C. et al. Combined inhibition of DDR1 and Notch signaling is a therapeutic strategy for KRAS-driven lung adenocarcinoma. Nat Med 22, 270–277, doi:10.1038/nm.4041 (2016).

56 Cespedes, M. V. et al. Selective depletion of metastatic stem cells as therapy for human colorectal cancer. EMBO Mol Med 10, doi:10.15252/emmm.201708772 (2018).

57 Gao, M. et al. Discovery and optimization of 3-(2-(Pyrazolo[1,5-a]pyrimidin-6-yl)ethynyl)benzamides as novel selective and orally bioavailable discoidin domain receptor 1 (DDR1) inhibitors. J Med Chem 56, 3281–3295, doi:10.1021/jm301824k (2013).

58 Geller, L. T. et al. Potential role of intratumor bacteria in mediating tumor resistance to the chemotherapeutic drug gemcitabine. Science 357, 1156–1160, doi:10.1126/science.aah5043 (2017).

59 Smith, T. J. et al. Recommendations for the Use of WBC Growth Factors: American Society of Clinical Oncology Clinical Practice Guideline Update. J Clin Oncol 33, 3199–3212, doi:10.1200/JCO.2015.62.3488 (2015).

60 Li, X. et al. Extracellular lumican inhibits pancreatic cancer cell growth and is associated with prolonged survival after surgery. Clin Cancer Res 20, 6529–6540, doi:10.1158/1078-0432.CCR-14-0970 (2014).

61 Kim, M. P. et al. Generation of orthotopic and heterotopic human pancreatic cancer xenografts in immunodeficient mice. Nat Protoc 4, 1670–1680, doi:10.1038/nprot.2009.171 (2009).

62 Oh, H., Siano, B. & Diamond, S. Neutrophil isolation protocol. J Vis Exp, doi:10.3791/745 (2008).

63 Chou, R. H. et al. EGFR modulates DNA synthesis and repair through Tyr phosphorylation of histone H4. Dev Cell 30, 224–237, doi:10.1016/j.devcel.2014.06.008 (2014).

64 Milne, T. A., Zhao, K. & Hess, J. L. Chromatin immunoprecipitation (ChIP) for analysis of histone modifications and chromatin-associated proteins. Methods Mol Biol 538, 409–423, doi:10.1007/978-1-59745-418-6_21 (2009).

