## Supplementary figure 1 for "DDR1-INDUCED NEUTROPHIL EXTRACELLULAR TRAPS DRIVE PANCREATIC CANCER METASTASIS"

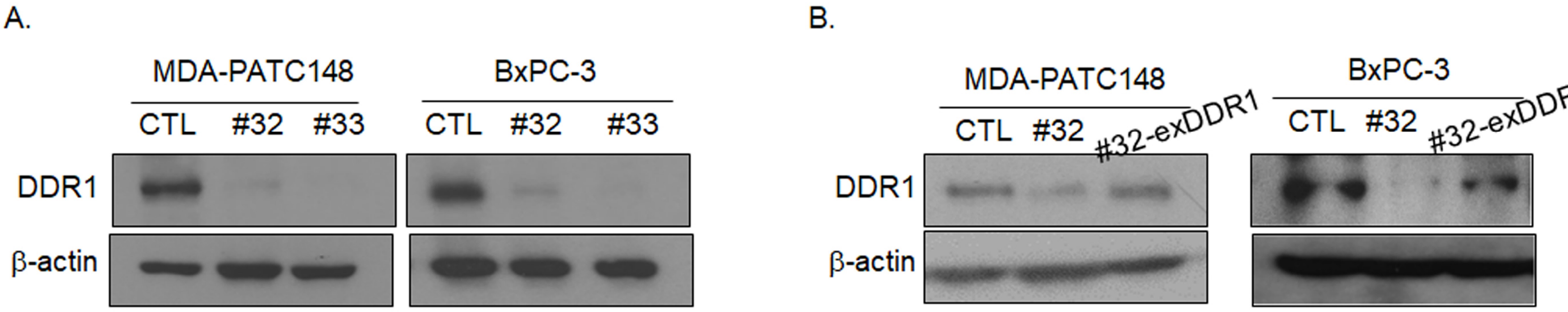


**Supplementary Figure 1.** DDR1 expression were analyzed by western blotting. **A** in MDA-PATC 148 and BxPC-3 cells with shRNA knockdown DDR1. **B** in MDA-PATC $148^{KD\#32}$ and BxPC-$3^{KD\#32}$ cells with re-express DDR1.
