## Supplementary figure 2 for "DDR1-INDUCED NEUTROPHIL EXTRACELLULAR TRAPS DRIVE PANCREATIC CANCER METASTASIS"

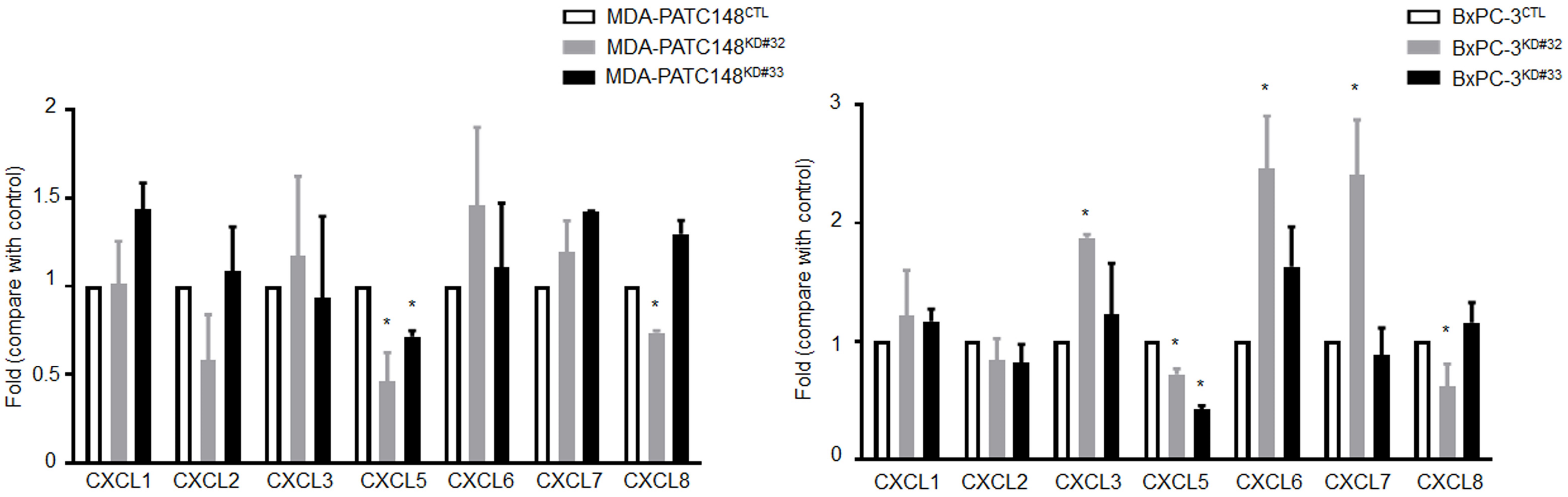


**Supplementary Figure 2.** DDR1 regulated CXCL5 mRNA level in both MDA-PATC 148 and BxPC-3 cells. Neutrophil-related chemokines mRNA level were analyzed by real-time PCR in MDA-PATC 148 and BxPC-3 cells with shRNA knockdown DDR1.
