## Supplementary figure 3 for "DDR1-INDUCED NEUTROPHIL EXTRACELLULAR TRAPS DRIVE PANCREATIC CANCER METASTASIS"

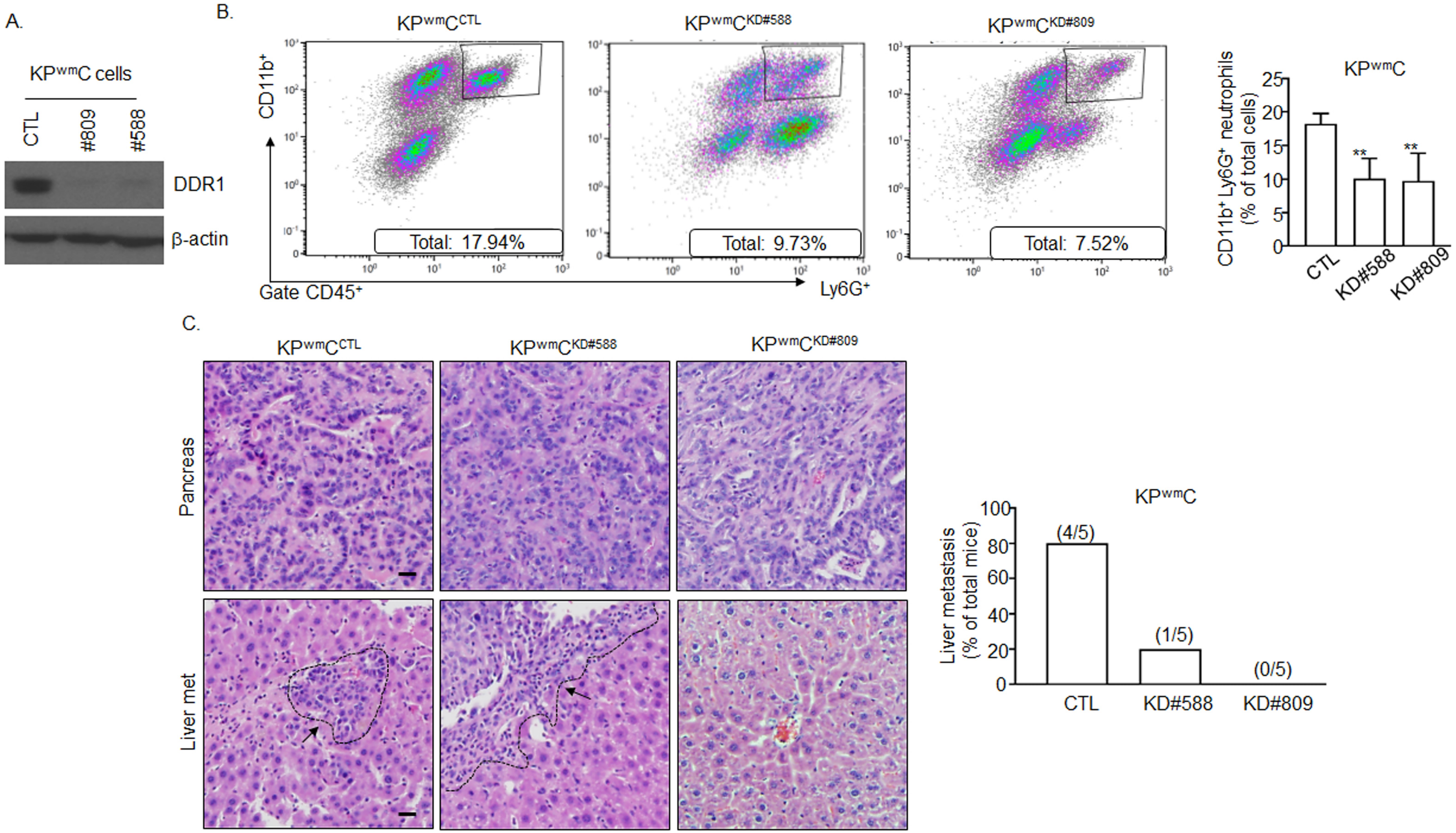


**Supplementary Figure 3.** DDR1 knockdown reduced CD11b+Ly6G neutrophils infiltration and liver metastasis. A DDR1 expression were analyzed by western blotting in KP$^{wm}$C cells with DDR1 knockdown. **B and C** C57BL/6J Mice were orthotopically injected with KP$^{wm}$C (control and 2 of DDR1-deficient clones) cells for 9 weeks. **B** *left:* CD11b+Ly6G+ neutrophils infiltrated into primary tumors were analyzed by FACS. *right:* the calculation of CD11b+Ly6G+ neutrophils infiltration were based on left panel (n=5 for each group). **C** *left:* H&E staining of pancreas and liver section. *Arrow*: region of tumor (5 mice per group), Scale bar, 50 µm. *right:* The numbers of liver-met calculation ($n = 5$ for each group).
