## Supplementary figure 4 for "DDR1-INDUCED NEUTROPHIL EXTRACELLULAR TRAPS DRIVE PANCREATIC CANCER METASTASIS"

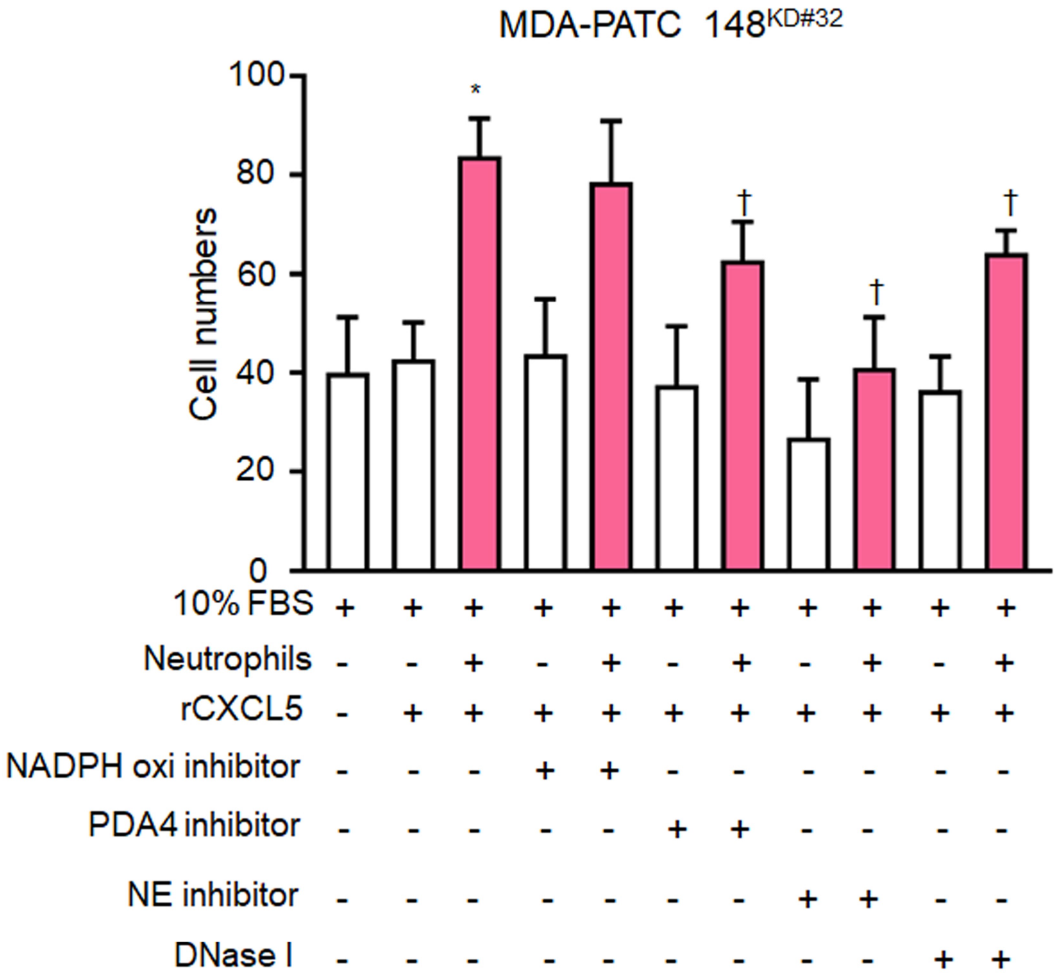


**Supplementary Figure 4.** CXCL5 mediated cancer cell invasion through NET. The invaded MDA-PATC 148KD#32 cells were analyzed by matrigel transwell chamber, co-culture with human neutrophils, with or without recombinant CXCL5, NDAPH oxidase, PDA4, NE inhibitor and DNase I treatment for 18 hours, and the average number of cells was calculated based on the number of cells found in six fields per chamber
