## Supplementary figure 5 for "DDR1-INDUCED NEUTROPHIL EXTRACELLULAR TRAPS DRIVE PANCREATIC CANCER METASTASIS"

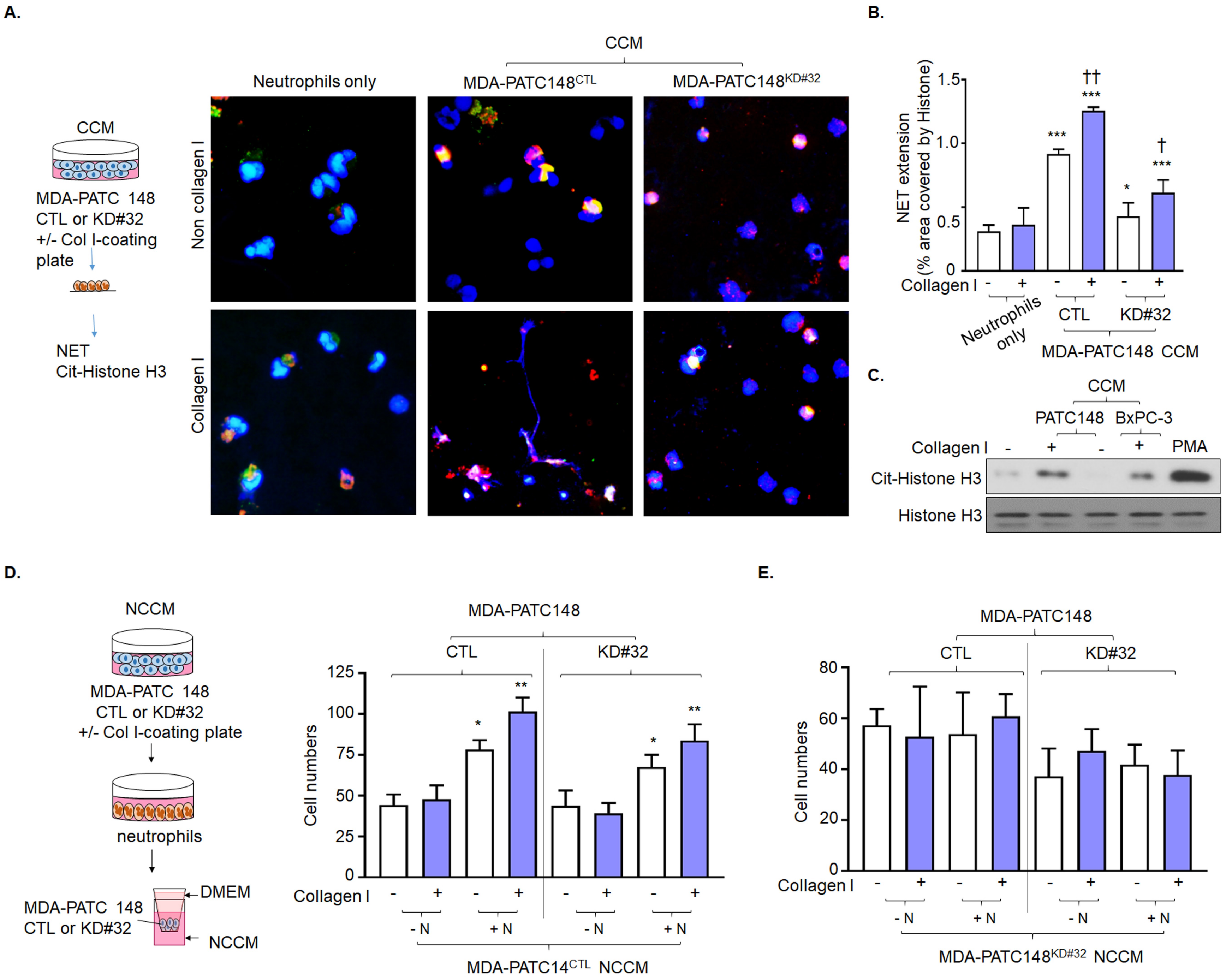


**Supplementary Figure 5.** DDR1-positive pancreatic cancer cells mediated NET formation from neutrophils through a soluble factor secretion and enhanced cancer cell invasion. **A-D** Human neutrophils were cultured with CCM from cancer cells for 16 hours. **A** NET structures were analyzed by immunofluorescence staining using DAPI (*blue*), anti-NE (*red*) and anti-histone (*green*) mAbs. *Scale bar*, 50 µm. **B** The NET quantification is displayed as NET histone area (µm$^2$) /per filed. **C** Cit-histone H3 expression were analyzed by western blotting. **D** The number of invaded cells analyzed by immunofluorescence staining using DAPI and calculated based on the number of cells found in six fields /per chamber.
