## Supplementary figure 6 for "DDR1-INDUCED NEUTROPHIL EXTRACELLULAR TRAPS DRIVE PANCREATIC CANCER METASTASIS"

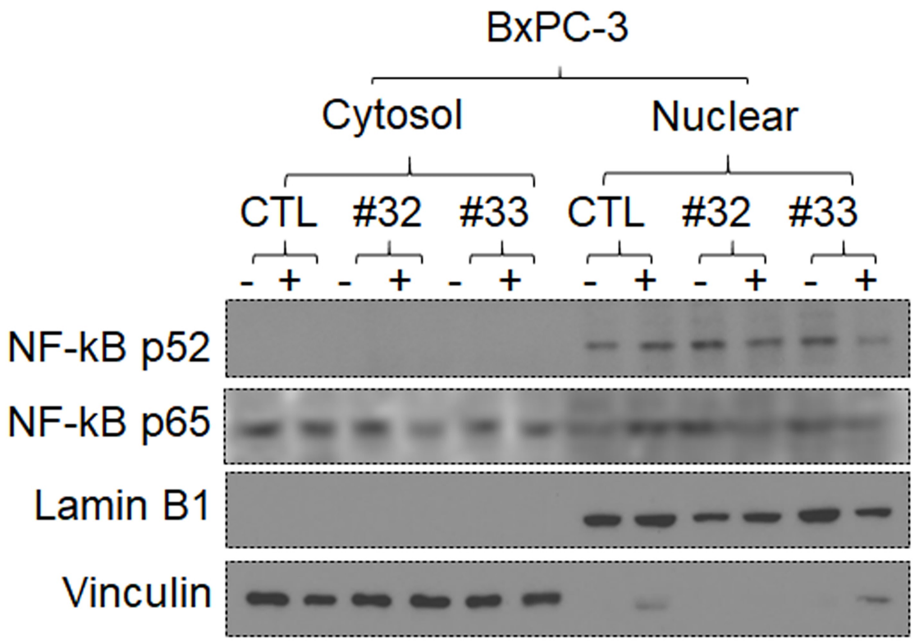


**Supplementary Figure 6.** DDR1 regulated NF-kB translocation into nucleus in BxPC-3 cells. Activated NF-kB were analyzed by western blotting using nuclear fraction in BxPC-3 cells with DDR1 knockdown.
