## Supplementary figure 7 for "DDR1-INDUCED NEUTROPHIL EXTRACELLULAR TRAPS DRIVE PANCREATIC CANCER METASTASIS"

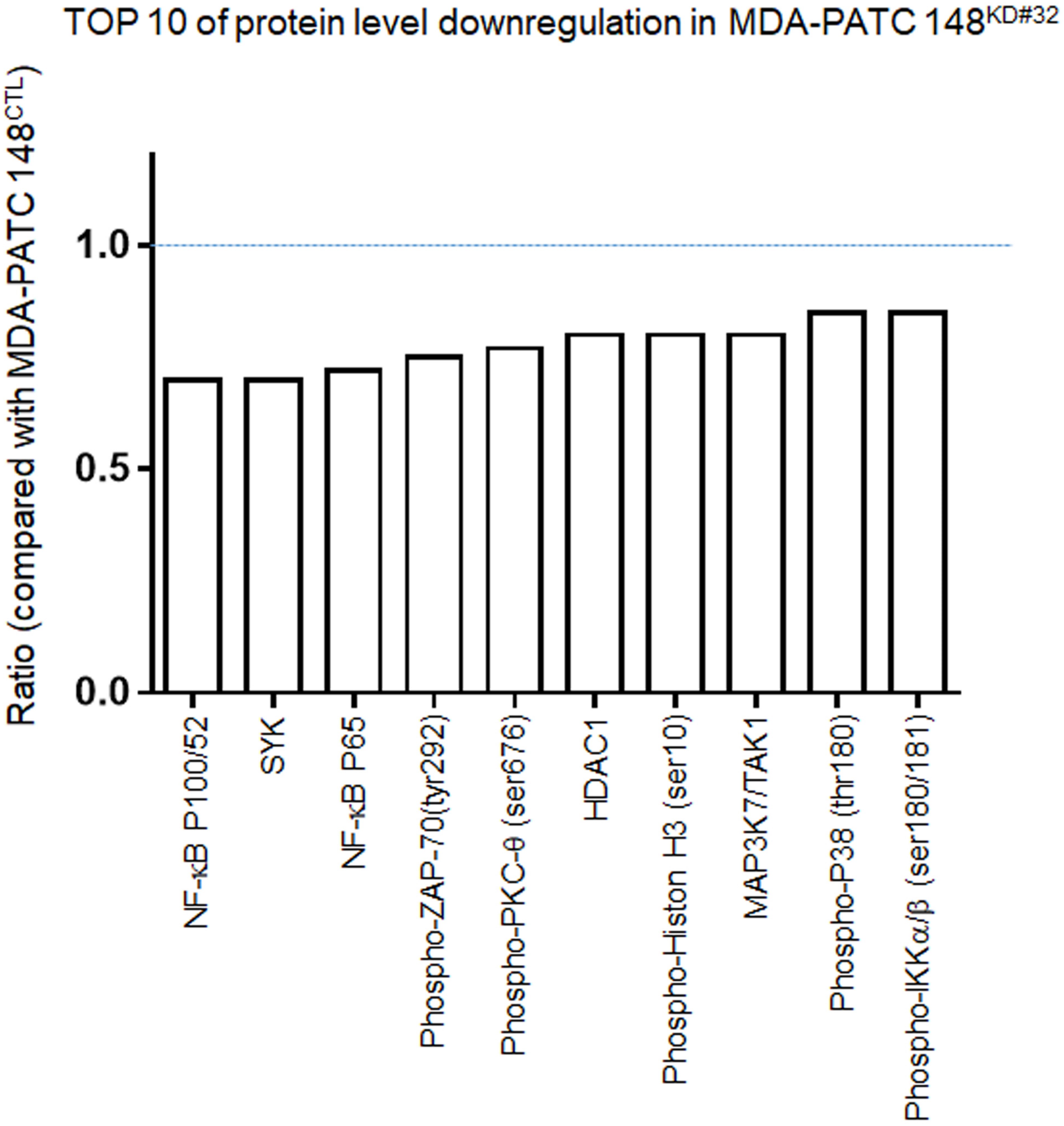


**Supplementary Figure** 7 Top 10 of protein level downregulation by using NFκB phospho antibody array in MDA-PATC 148 cells with DDR1 knockdown.
