## Supplementary figure 8 for "DDR1-INDUCED NEUTROPHIL EXTRACELLULAR TRAPS DRIVE PANCREATIC CANCER METASTASIS"

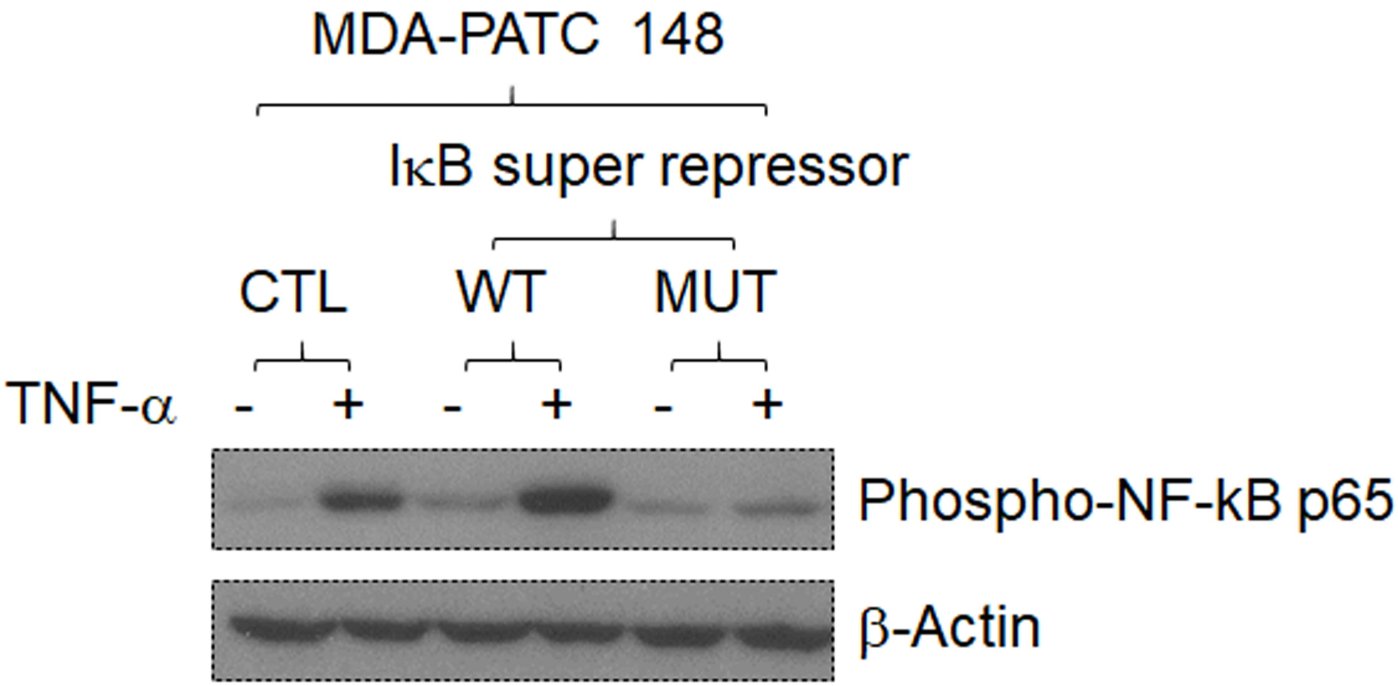


**Supplementary Figure 8.** Activated NF-kB P65 were dececected by western blotting in MDA-PATC 148 with IκB super-repressor mutation.
