## Supplementary figure 9 for "DDR1-INDUCED NEUTROPHIL EXTRACELLULAR TRAPS DRIVE PANCREATIC CANCER METASTASIS"

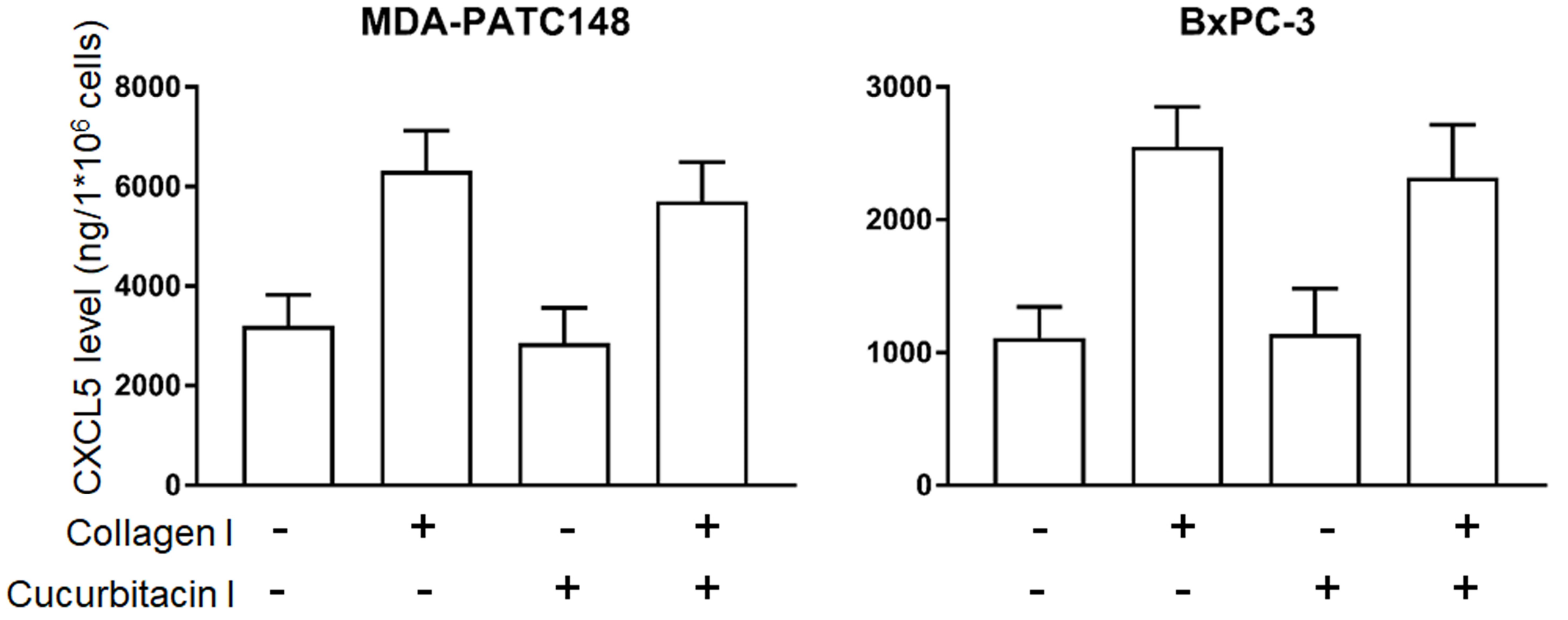


**Supplementary Figure 9** Collagen I induced CXCL5 level through STAT3-independent pathway. CXCL5 level were analyzed by ELISA in MDA-PATC 148 and BxPC-3 cells with cucurbitacin I pretreatment for 30 mins, with or without collagen I treatment for 3 hours.
