## Supplementary figure 10 for "DDR1-INDUCED NEUTROPHIL EXTRACELLULAR TRAPS DRIVE PANCREATIC CANCER METASTASIS"

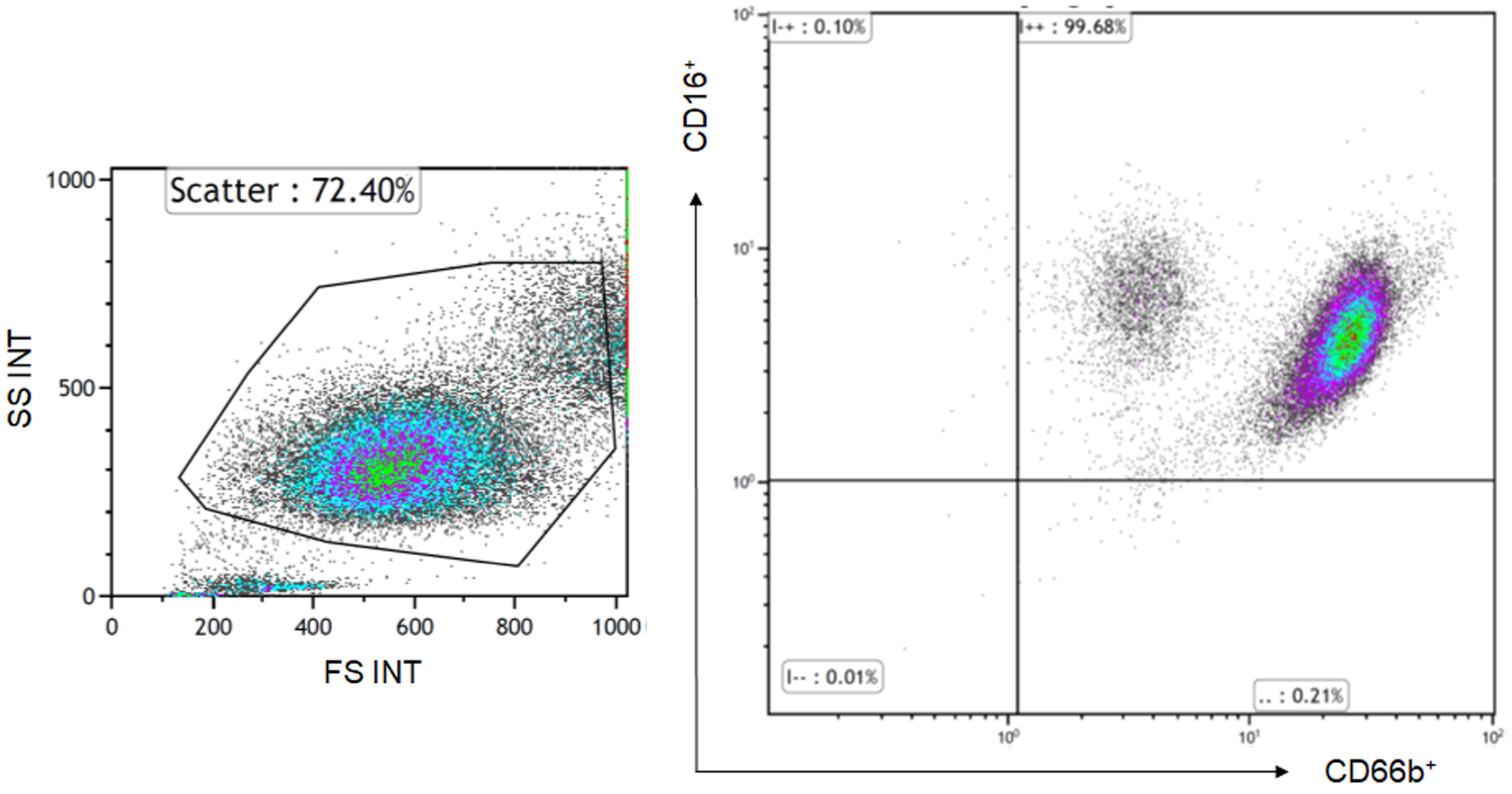


**Supplementary Figure 10.** Human neutrophils from human blood were checked by FACS using anti-CD16 and anti-CD66b antibodies.
