## Supplementary figure 11 for "DDR1-INDUCED NEUTROPHIL EXTRACELLULAR TRAPS DRIVE PANCREATIC CANCER METASTASIS"

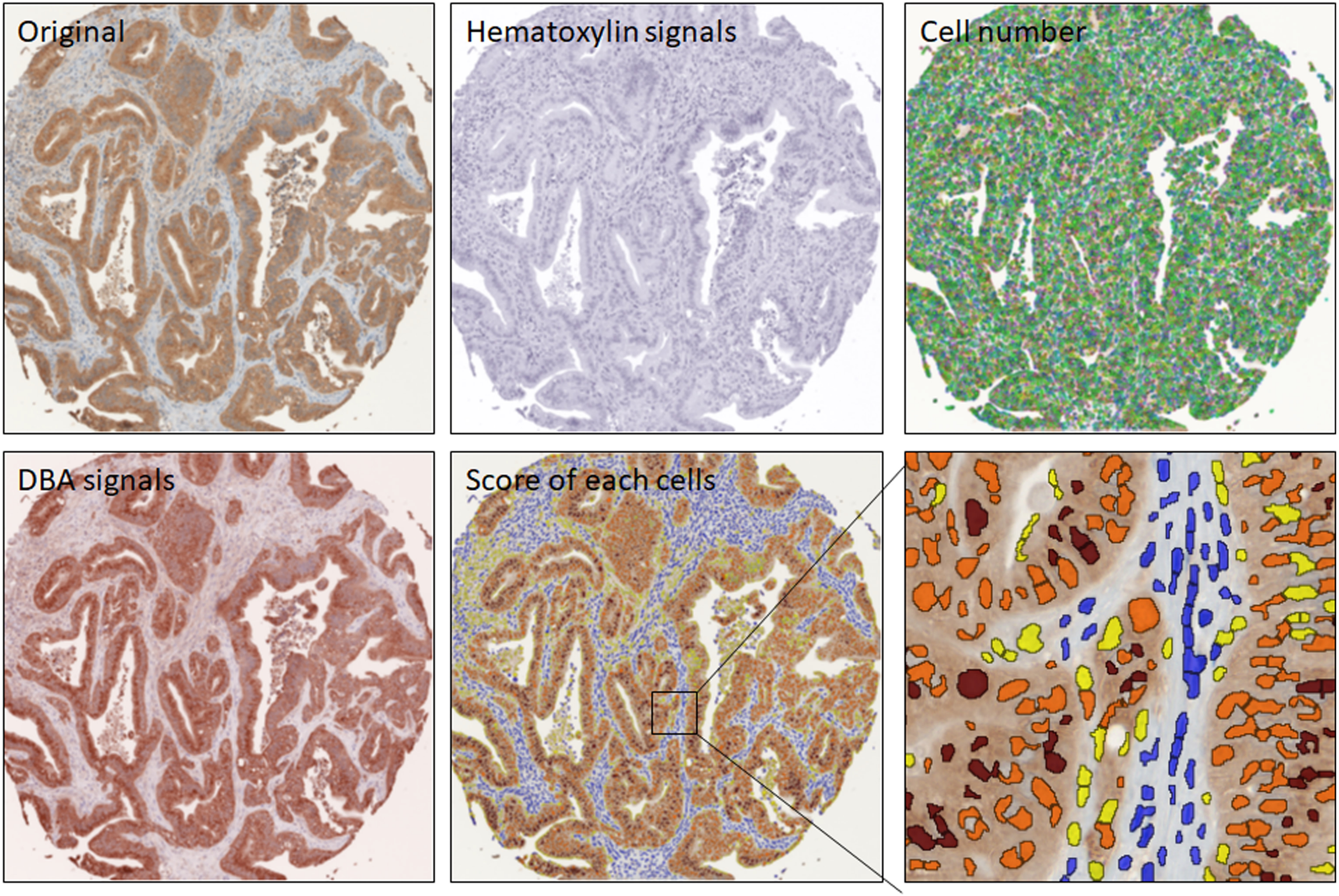


**Supplementary Figure 11.** Quantification of IHC signal. Images were scanned using PE Vectra3 and processed by inform[R] software, the H-Score were calculated by DBA signals/per selected cells. The score of each cells is 0 showed in blue, 1 showed in yellow, 2 showed in orange and 3 showed in brown.
