## Supplementary figure 12 for "DDR1-INDUCED NEUTROPHIL EXTRACELLULAR TRAPS DRIVE PANCREATIC CANCER METASTASIS"

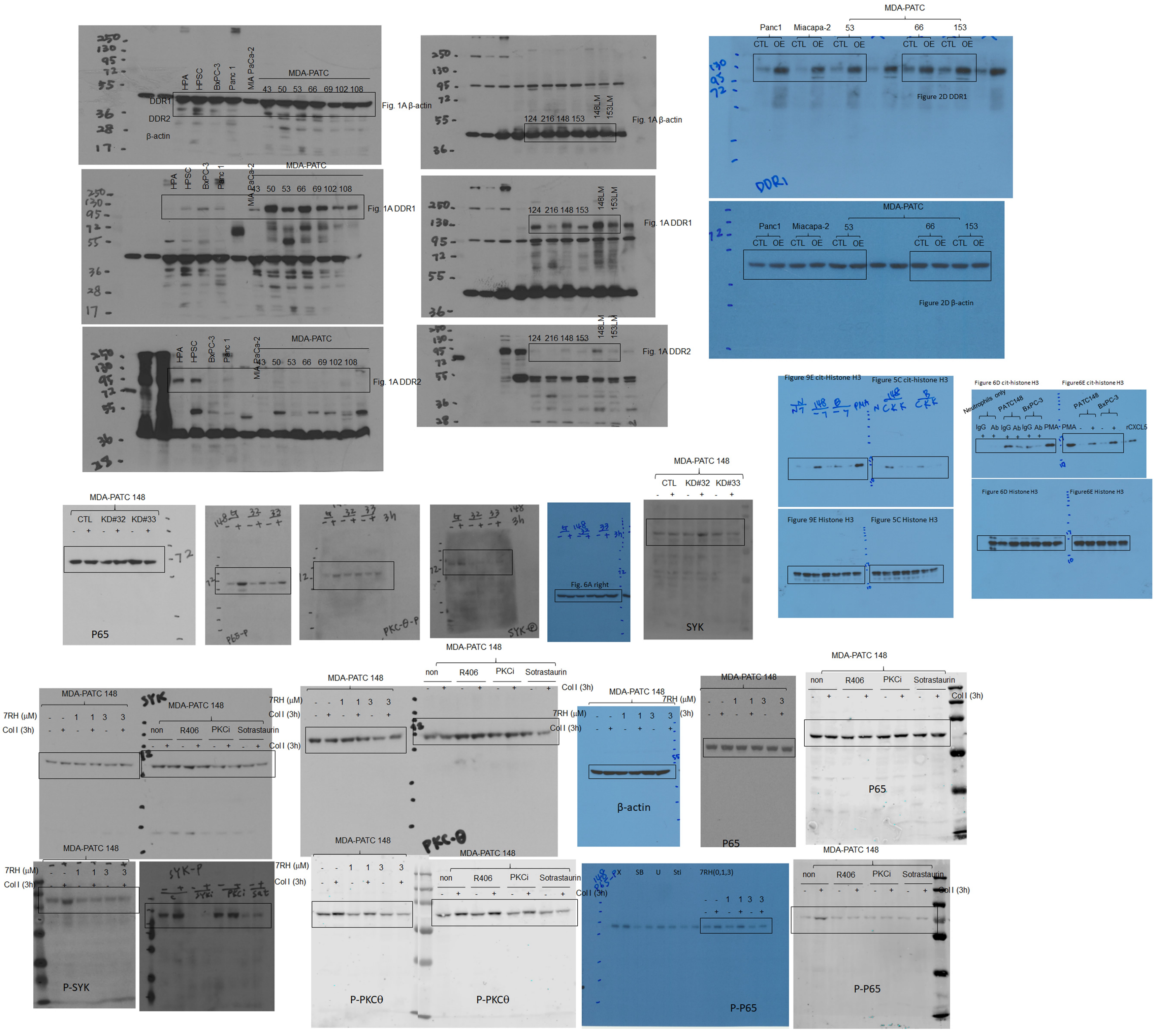


**Supplementary Figure 12.** Original lumiescence images of the blots presented in the main text . Related to Figure 1-9.
