## Supplementary table 1 for "DDR1-INDUCED NEUTROPHIL EXTRACELLULAR TRAPS DRIVE PANCREATIC CANCER METASTASIS"

| Primer for real-time PCR | Sequence 5' to 3' |
|---|---|
| hCXCL1-F | GAAAGCTTGCCTCAATCCTG |
| hCXCL1-R | CTTCCTCCTCCCTTCTGGTC |
| hCXCL2-F | GGGCAGAAAGCTTGTCTCAA |
| hCXCL2-R | GCTTCCTCCTTCCTTCTGGT |
| hCXCL3-F | CGCCCAAACCGAAGTCATAG |
| hCXCL3-R | GCTCCCCTTGTTCAGTATCTTTT |
| hCXCL5-F | TGGACGGTGGAAACAAGG |
| hCXCL5-R | CTTCCCTGGGTTCAGAGAC |
| hCXCL6-F | AGAGCTGCGTTGCACTTGTT |
| hCXCL6-R | GCAGTTTACCAATCGTTTTGGGG |
| hCXCL7-F | GGCTTCCTCCACCAAAGGAC |
| hCXCL7-R | TCTTTGCCTTTCGCCAAGTT |
| hCXCL8-F | GAATGGGTTTGCTAGAATGTGATA |
| hCXCL8-R | CAGACTAGGGTTGCCAGATTTAAC |
| hGAPDH-F | ACGGATTTGGTCGTATTGGG |
| h-GAPDH-R | TGATTTTGGAGGGATCTCGC |
| Primer for ChIP assay | |
| CXCL5 promoter-NFκB-f-1 | TAGAGGTGCACGCAGCTCCT |
| CXCL5 promoter-NFκB-r-1 | GAGCACTGTGGCTTCCTCGT |
